# CUBIC-Cloud: An Integrative Computational Framework Towards Community-driven Whole-Mouse-Brain Mapping

**DOI:** 10.1101/2020.08.28.271031

**Authors:** Tomoyuki Mano, Ken Murata, Kazuhiro Kon, Chika Shimizu, Hiroaki Ono, Shoi Shi, Rikuhiro G. Yamada, Kazunari Miyamichi, Etsuo A. Susaki, Kazushige Touhara, Hiroki R. Ueda

## Abstract

Recent advancements in tissue clearing technologies have offered unparalleled opportunities for researchers to explore the whole mouse brain at cellular resolution. With the expansion of this experimental technique, however, a scalable and easy-to-use computational tool is in demand to effectively analyze and integrate whole-brain mapping datasets. To that end, here we present CUBIC-Cloud, a cloud-based framework to quantify, visualize and integrate whole mouse brain data. CUBIC-Cloud is a fully automated system where users can upload their whole-brain data, run analysis and publish the results. We demonstrate the generality of CUBIC-Cloud by a variety of applications. First, we investigated brain-wide distribution of PV, Sst, ChAT, Th and Iba1 expressing cells. Second, Aβ plaque deposition in AD model mouse brains were quantified. Third, we reconstructed neuronal activity profile under LPS-induced inflammation by c-Fos immunostaining. Last, we show brain-wide connectivity mapping by pseudo-typed Rabies virus. Together, CUBIC-Cloud provides an integrative platform to advance scalable and collaborative whole-brain mapping.

## Introduction

Massive and collective observation of complex systems (often referred to as omics approaches) is the driving force of modern biology. It would also be true for studying highly-evolved mammalian brains, where a complex system of intricately connected cells gives rise to intelligent behaviors. In particular, comprehensive approaches to identifying the properties of every single cells *in situ* within this complex system would be pivotal. Recent advancements in tissue clearing technology have brought new breakthroughs in this landscape [1, 2, 3, 4, 5, 6, 7, 8, 9]. Combined with light-sheet fluorescence microscopy (LSFM) [10] and genetical, viral and immunohistochemical labelling techniques, tissue clearing now enables high-speed volumetric imaging of mammalian (most prominently mouse) brains at cellular resolution [7, 11, 12, 13, 14, 15, 16, 17]. Built on top of these technological advancements, we recently reported the construction of CUBIC-Atlas [18], a 3D mouse brain atlas with single cell resolution, where all of the cells in the brain (amounting to approximately 0.1 billion) were digitally analyzed and recorded.

These scientific advancements encourage us to conceive a future where whole-brain mapping projects, which conventionally required institution-scale resources and efforts, can be carried out by individual laboratories, or even by a single researcher [19, 20, 21, 22]. In this regard, the current technological stage can be thought of as parallel to the dawn of genome sequencing technology in early 2000s. In genome science, the importance of the data repository cannot be understated. The emergence of the database to browse and search genomes (such as UCSC Genome Browser [23]) played a critical role in integrating data collected in numerous sites across the globe. Such distributed collaboration prompted a rapidly growing coverage of various organisms and individuals, pioneering the data-driven discoveries of gene functions and new therapeutics. In neuroscience, several large-scale mouse brain datasets have been constructed, such as Allen Mouse Brain Atlas [24] and Brain Architecture Project [25], primarily using serial sectioning tomography methods. However, a common platform has been yet to appear that offers the opportunity for the community to submit and share new data, embracing the tissue clearing and rapid brain scanning techniques. Based on these considerations, we suggest that it is now possible to construct the community-supported mouse brain data repository, borrowing collaborative ideas from genome sciences.

By referring to the previous image analysis pipelines for tissue clearing samples [26, 27, 28, 16, 29], we postulated that the following elements should be considered in composing a framework for whole mouse brain mapping. First, the reference brain (equivalent to the template sequence), to which all brain data are aligned, is necessary. We think that CUBIC-Atlas would play a central role in addressing this challenge. Second, the framework should be constructed around the research community, which allows researchers to submit the data, as well as to browse and search the brains in the previous studies. Third, because of the complexity and the large size of the whole-brain data sets, the framework should be equipped with a toolkit to visualize and quantify the data to assist intuitive understanding. Fourth, these software tools should offer superior accessibility and usability to the end users, without requiring specialized expertise in programming or powerful computer resources.

In this paper, we present a computational framework for single-cell-resolution whole-mouse-brain analysis, named CUBIC-Cloud. We built CUBIC-Cloud on top of the latest cloud computing technologies. After identifying single cells from 3D image stack, users can upload the structure image and detected cell list to the CUBIC-Cloud server. CUBIC-Cloud server automatically aligns individual brains with the CUBIC-Atlas, and construct the user’s own mouse brain database. CUBIC-Cloud also offers GUI tools to perform various kinds of quantification tasks. Uploaded brains can be interactively visualized using 3D brain viewer. The scientific results obtained thereby can be easily published in the CUBIC-Cloud’s public repository, to allow other researchers to view the data. CUBIC-Cloud is hosted at https://cubic-cloud.com.

After describing the software architecture, we extensively demonstrate the capability and generality of CUBIC-Cloud framework by analysing over 50 whole mouse brains, covering four important application domains: (1) investigating the distribution of targeted cell types (“cell-type mapping”), (2) quantifying the pathological state by disease markers, (3) reconstructing the neuronal activity profile by imaging the expression of c-Fos (“activity mapping”), and (4) deciphering brain-wide connectivity using Rabies virus tracers (“circuit mapping”). With these applications, we established a general framework to effectively integrate whole-brain mapping experiments, offering new opportunities for data-driven discoveries in neuroscience.

## Results

### CUBIC-Cloud whole-brain analysis framework

To design a unified workflow for whole-mouse-brain analysis, we constructed CUBIC-Cloud on top of our previously published tissue clearing protocols and brain mapping strategies [7, 18, 30]. The workflow of whole-brain analysis using CUBIC-Cloud is illustrated in Fig. 1. Briefly, the workflow divides into (1) tissue clearing and image collection, (2) single cell detection, and (3) uploading the data to CUBIC-Cloud, where brain registration, brain-wide quantification and visualization are performed.

**Fig. 1.**
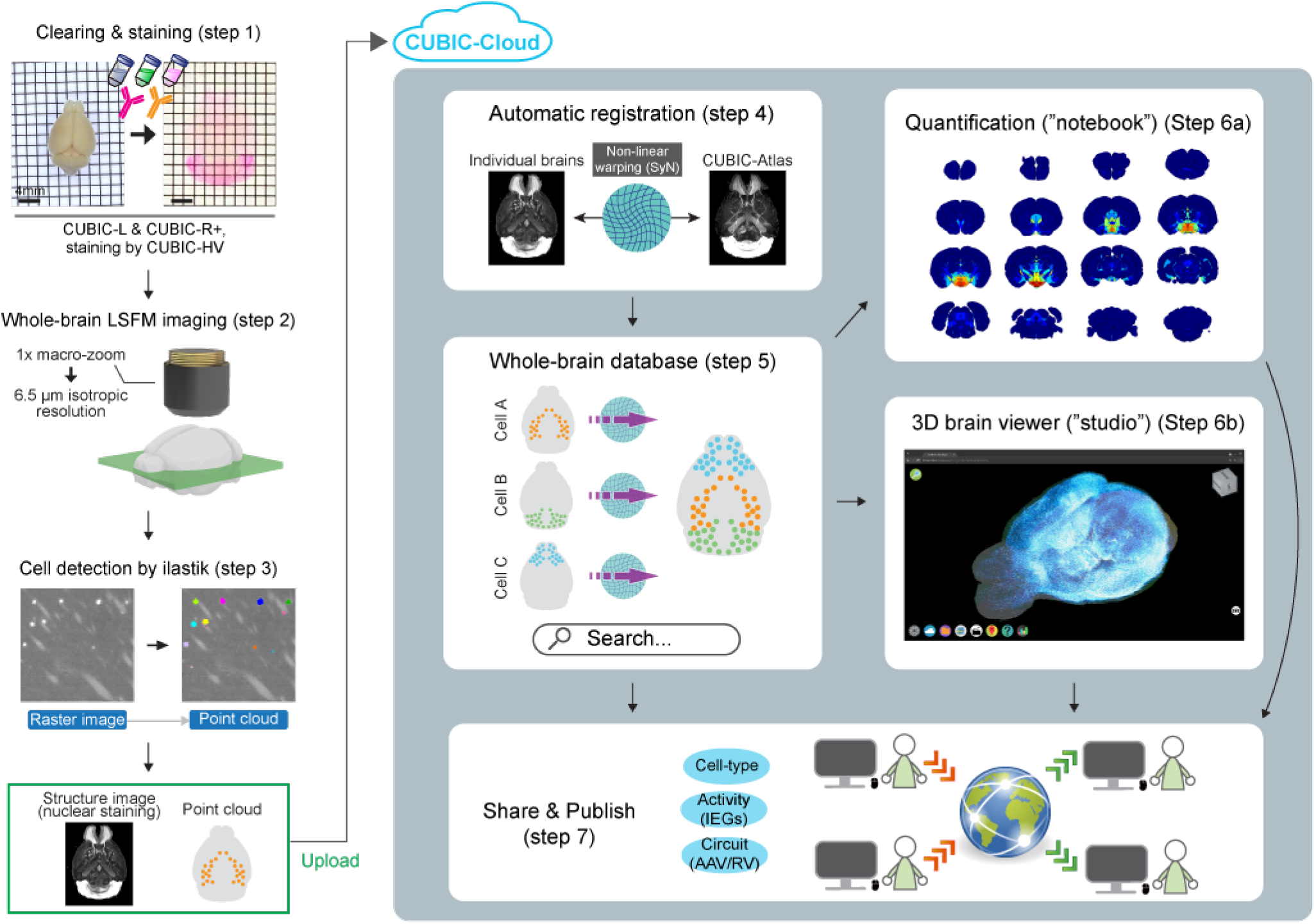
CUBIC-Cloud: A Cloud-based Computational Framework for Whole Mouse Brain Analysis. The overview of whole-brain analysis pipeline by CUBIC-Cloud. In this study, mouse brains were cleared by CUBIC-L and CUBIC-R+ reagents and 3D-stained by CUBIC-HV protocol. Cleared brains were imaged using macro-zoom LSFM. From the obtained image stacks, single cells were isolated using ilastik software, converting the raw raster image into an ensemble of discrete cells (i.e. point cloud). Users then upload the point cloud cells and structure image to CUBIC-Cloud. In the cloud, brain registration is automatically performed to align individual brains with the reference brain. Thereby, user’s own brain database is constructed. Then, user can perform various kinds of brain-wide quantification using “notebook”. CUBIC-Cloud also offers an interactive 3D whole-brain viewer (“studio”). Lastly, CUBIC-Cloud lets the user share and publish their point-cloud whole-brain data, as well as notebooks and studios, to allow broad access from the researchers.

In this study, mouse brains were cleared by CUBIC-L and CUBIC-R+ reagents [30] (“step 1” in Fig. 1). In addition, nuclear counter-staining (for brain registration) and, optionally, immunostaining were applied by following the CUBIC-HV protocol [31] (Methods). In most experiments, cleared brains were imaged using macro-zoom LSFM with 6.5 μm isotropic XYZ voxel size (“step 2” in Fig. 1, Methods). The transparency of the cleared brain was quantitatively evaluated by imaging fluorescent beads embedded in the tissue, validating homogeneous image quality throughout all brain regions (Supplementary Fig. 2, Methods). Of note, CUBIC-Cloud framework should be general enough to be used with other clearing methods, as long as the equivalent level of tissue transparency and morphology preservation is achieved.

After obtaining the 3D image stack, labeled cells were detected using ilastik software [32, 33] (“step 3” in Fig. 1). ilastik uses supervised machine learning (random forest algorithm) to classify pixels into multiple classes (such as cells and backgrounds), which performed robustly well even when cells from different brain regions presented diverse brightness and morphology. We created a Python script which wrapped ilastik to automate the workflow and add some custom analysis routines (Methods). The accuracy of cell counting is described in the corresponding sections. By this cell counting procedure, raw raster images were converted into a discrete ensemble of cellular points, or point cloud, where positions, volumes and fluorescence intensities of all labeled cells in the brain were recorded. Of note, users can opt to use their own cell detection routine, as long as the output follows the required format by CUBIC-Cloud.

After cell detection, the rest of the analysis were carried out by CUBIC-Cloud. Users can go to the CUBIC-Cloud web site (https://cubic-cioud.com) and upload cellular point cloud and corresponding structure image (nuclear staining channel) to the server. CUBIC-Cloud is deployed on Amazon Web Service (AWS) using so-called serverless architecture. Serverless architecture allows the cloud to dynamically scale its capacity based on the computational load, enabling the cloud to handle practically unlimited number of tasks in parallel while minimizing the idling time. The detailed implementation is described in Methods (also see Supplementary Fig. 1). Once the brain data is uploaded to the CUBIC-Cloud server, it is automatically sent to the preprocessing task queue, which is powered by ECS and EC2 on AWS. Preprocessing task first performs brain registration to align submitted brain with the CUBIC-Atlas (“step 4” in Fig. 1). CUBIC-Cloud uses the symmetric diffeomorphic image registration (SyN) algorithm implemented in ANTs library [34]. Using the nuclear staining channel, SyN iteratively optimized the warp field to maximize the normalized cross-correlation (NCC), which measured the similarity between two images (Supplementary Fig. 3i,j, Methods). After registration, cell coordinates are transformed to the CUBIC-Atlas space, and all cells are given anatomical region IDs (following the Allen Brain Atlas CCF v3 [24]).

By repeating the above procedures, users can construct their own brain database in CUBIC-Cloud. As the database grows, users can search brains using tags attached to them, such as cell labels and project names, along with other metadata. To perform quantitative analysis of the brains in the database, CUBIC-Cloud offers a “notebook”, a feature that allows users to create various kinds of graphs with graphical user interfaces (GUIs) (“step 6a” in Fig. 1). To give some concrete examples, many of the figure items presented in this paper were created by CUBIC-Cloud’s notebook.

CUBIC-Cloud also offers an interactive 3D brain viewer, a feature called “studio” (“step 6b” in Fig. 1). Here, brain data is visualized in point cloud format, where the whole brain is rendered as an ensemble of discrete cellular points, each carrying biological attributes such as protein expression levels. Point cloud format is more efficient to deliver data to the remote clients than sending raw raster data, while carrying the essential biological information. The viewer runs on standard web browsers, and the point clouds are adaptively queried from the server upon the viewer’s camera movement, enabling to interact with millions of cells in real time.

The last essential component of CUBIC-Cloud is sharing and publishing. Users can share their whole-brain data with specific users, such as research collaborators, and grant access to the data. Users can opt to publish their data in the CUBIC-Cloud’s public repository. Once published, any users can view the brain. The share and publish capability are also supported for the notebooks and studios. Therefore, users can transparently show their analysis results to the research community. To demonstrate this concept, all of the brain data investigated in this study is deposited on CUBIC-Cloud public repository, as well as the notebooks and studios that performed the analysis. Together, CUBIC-Cloud offers cloud-native and integrative software solution for whole-mouse-brain mapping.

### Whole-brain analysis of PV, SST, ChAT and Th expressing cells

In the following, we use CUBIC-Cloud in several neuroscience applications to show the generality of the proposed framework. As the first application, we attempted to quantify the whole-brain distribution of distinct cellular subtypes that express the following markers, respectively: parvalbumin (PV), somatostatin (SST), choline acetyltransferase (ChAT), tyrosine hydroxylase (TH), and ionized calcium-binding adapter molecule 1 (Iba1). Whole-brain scale analysis of PV, SST and ChAT expressing neurons were previously reported using knock-in transgenic mouse line with Cre recombinase and fluorescent proteins [25, 35, 36]. However, no such whole-brain scale analysis have been done so far using immunostaining and tissue clearing. Immunostaining offers several orthogonal advantages over genetical approaches, such as the capability to assess the absolute expression amount, and thus of significant value. Therefore, we used CUBIC-HV protocol [31] to label the whole adult brain tissue with PV, SST, ChAT, TH and Iba1 antibody, respectively (Methods).

The obtained images contained strong and soma-localized signals as well as moderately bright fibrous structures. Here, we attempted to quantify the number of immunoreactive cell bodies, and thus, the ilastik classifier was trained to isolate cell soma and reject fiber-like signals. The accuracy of the cell detection was extensively evaluated (Supplementary Fig. 3c-g). Our cell detection demonstrated 75-95% precision and 60-85% sensitivity for the regions evaluated, which we concluded was accurate enough to perform quantitative analysis.

Whole-brain views of the investigated cell types are shown in Fig. 2a-f (rendered by CUBIC-Cloud’s 3D viewer). Here, cells are psuedo-colored, reflecting the fluorescence intensity of the immunostaining. In our analysis, the total number of detected cells were: (6.1 ±0.7) × 10^5^ (PV, *n* = 4); (6.7±0.3) × 10^5^ (SST, *n* = 4); (6.5±0.2) × 10^4^ (ChAT, *n* = 4); (6.9± 1.2) × 10^4^ (TH, *n* = 4); (2.72±0.14) × 10^6^ (Iba1, *n* = 7) (Suplementary Table 2, Suplementary Table 3). In this paper, mean ± standard deviation is used unless otherwise specified. The relative ratio of each cell type in all brain regions except isocortex is shown in Fig. 2g. Here, the ratio was calculated by dividing the detected cell count by the total number of cells reported in CUBIC-Atlas [18]. We will discuss the brain-wide distribution of these cell types in more detail in Methods.

**Fig. 2.**
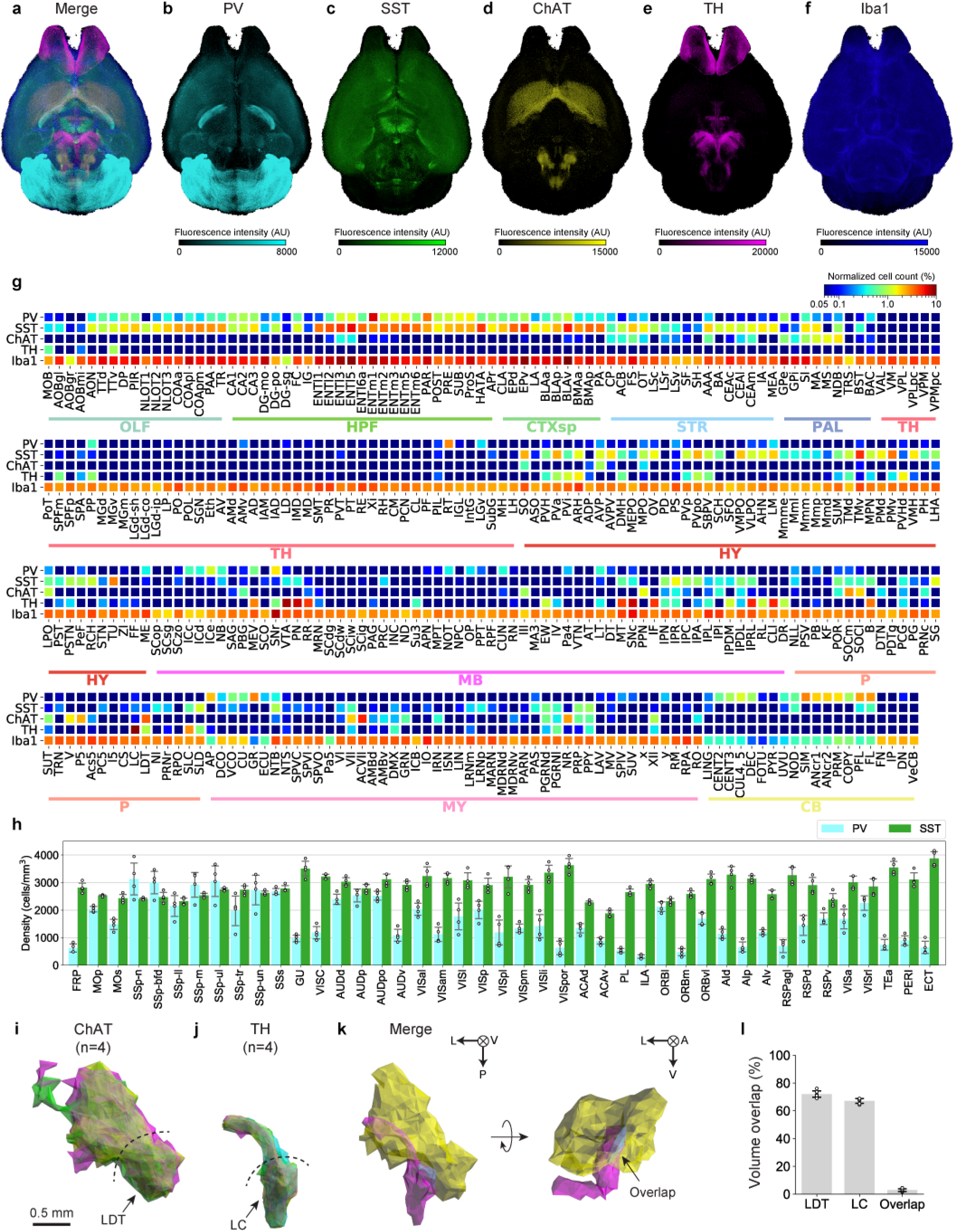
Whole-brain Analysis of PV, Sst, ChAT and Th Expressing Cells. Whole-brain distribution of PV, SST, ChAT, TH and Iba1 expressing cells were investigated by applying 3D immunostaining and by using CUBIC-Cloud analysis framework. **a-f**, Whole-brain views of labelled cells. Each point (i.e. single cell) was assigned a pseudo-color based on its fluorescence intensity. **g**, Relative population heatmap of all brain regions outside the isocortex. The number of each cell type was normalized by the total number of cells in each regions, derived from CUBIC-Atlas (*n* = 4 for PV, SST, ChAT and TH; *n* = 7 for Iba1). **h**, Density of PV and Sst expressing cells in the isocortex. Data are shown as mean ± STD (*n* = 4). **i**, From the cluster of ChAT expressing cells, the boundary surface including LDT was extracted (*n* = 4). **j**, From the cluster of TH expressing cells, the boundary surface including LC was extracted (*n* = 4). **k**, Merge of the two boundary surfaces (yellow = ChAT, magenta = TH, cyan = overlapping region). **l**, The volume overlaps of the boudary surfaces. Brain region acronyms follow the ontology defined by the Allen Brain Atlas.

Within the isocortex, PV neurons were most densely populated in somatosensory and auditory areas, whereas there were relatively sparse populations in medial frontal and lateral association areas (Fig. 2h). Compared to PV, SST neurons were more homogeneously observed in the isocortex, consistent with the Cre-based study by [25]. Laminar density analysis revealed that there were almost no PV cells in layer 1, and that cell density reaches its peak in layer 4 or 5 for both PV and SST (Supplementary Fig. 4a-d), successfully recapitulating the previous observations [37, 38]. There were sparse populations of ChAT neurons, whose ChAT expression levels were very low (Supplementary Fig. 4e,f). These ChAT neurons were most dense in layer 2/3 or 4, corroborating the previous observation [37]. For Th neurons, no significant population above false-positive detection level were found in the isocortex. Iba1 expressing cells were ubiquitously observed in the isocortex, with similar density across layers.

From our cell-type mapping data, certain cell types were clustered in specific areas in the brain, offering the clues to delineate the region boundaries. For instance, we observed that ChAT neurons were densely populated in and around the LDT (for the list of region acronyms, see Suplementary Table 1). From this cluster of ChAT neurons, we defined the polygonal boundary enclosing these cells using alpha shape algorithm (Fig. 2i, Methods). We performed same analysis targeting TH neurons in and around the LC (Fig. 2j). The polygonal boundaries defined by ChAT+ and TH+ neuron cluster, respectively, were neighboring with each other with a small overlap (Fig. 2k). We then evaluated the overlap between polygonal boundaries using the Dice’s metric (Methods). The result showed about 70% overlap between ChAT-ChAT and TH-TH pairs (Fig. 2l). Furthermore, the overlap of ChAT-TH pair was 2.84±0.9 %. These results support the accuracy of our registration method with an orthogonal evaluation metric, which is largely independent from information present in the nuclear staining channel. This result thus offers the possibility to accurately delineate the brain regions by collecting more cell-type mapping data in CUBIC-Cloud.

Lastly, to assess whether our analysis pipeline can accurately quantify the expression amount of the proteins, we evaluated the expression of Iba1 under artificially induced inflammation state, because Iba1 expression is known to be correlated with microglial activation [39]. In our experiment, 1 mg/kg of lipopolysaccharides (LPS), a purified extract of the outer membrane of Gram-negative bacteria, was administered to mice via intraperitoneal (i.p.) injection. The brains were sampled twenty-four hours after injection (Methods). In the isocortex, no subregions displayed significant change in the cell density (*p* > 0.1, Welch’s t-test, Supplementary Fig. 4i), while the mean expression amount per cell was increased significantly in all subregions (*p* < 0.05, Welch’s t-test, Supplementary Fig. 4j). The observation that cortical microglia does not proliferate but increases its Iba1 expression upon LPS administration was reported in [40], and our results confirmed that it is globally true in all cortical areas. Outside the isocortex, most regions showed increases in Iba1 expression amount (Supplementary Fig. 4k), with varying degree of change. For instance, Iba1 expression amount barely changed in cerebellum. On the other hand, we confirmed that increase in Iba1 expression was markedly high in the SFO and IO, which are part of ircumventricular organs (CVOs) having highly permeable blood-brain barrier (BBB) [41].

### Whole-brain analysis of Aβ plaque accumulation in AD model mouse brain

We next applied CUBIC-Cloud analysis framework to quantitatively understand the pathological state of the Alzheimer’s disease (AD) model mouse. To demonstrate this, the whole brain from an *App*^NL-G-F/NL-G-F^ AD model mouse [42] (9-to 10-months-old) was cleared and stained by anti-Aβ antibody (*n* = 4, Methods). In LSFM images, Aβ plaques were observed as dim blobs often accompanying a bright spot at the core. On the other hand, no plaque staining pattern was observed in the control wild-type mouse brain (9-to 10-month-old, *n* = 3, data not shown).

We first quantified the density (number of individual plaques per volume) and the volume ratio (computed as (total plaque volume in the region)/(region volume)). In both metrics, Aβ plaque amounts were highest in the cerebral cortex and cereberal nuclei, and relatively lower amount of plaques were observed in the brain stem and cerebellum (Fig. 3a,b). The effective radius of the plaque (computed as *r* = {3*V*/(4***π***)}^1/3^ where *V* is the plaque volume) tended to be larger in the isocortex and hippocampus and smaller in cerebellum (Fig. 3c). Within the isocortex, a relatively stronger accumulation of Aβ were observed in visual and auditory areas, whereas plaques were relatively sparse in in medial frontal areas (Fig. 3d,e). Layer-wise abundance of Aβ plaque showed concave profile, with its peak in layer 4 (Fig. 3f). The whole-brain cartoon heatmap showing the Aβ volume ratio is shown in Fig. 3g. In the brain stem, the plaque volume ratio was typically 0.5% to 1.0%. Some brain stem regions, however, showed notably larger or smaller amount of Aβ accumulation. For example, SNr and VMH had relatively higher amount of Aβ compared to neighboring regions (Fig. 3h,i). ARH, right next to VMH, had almost no Aβ plaques. We also observed that the regions around the ventricles showed relatively lower amount of plaques, including TRS, DTN, MH, LH and PVT (Fig. 3g, j-l).

**Fig. 3.**
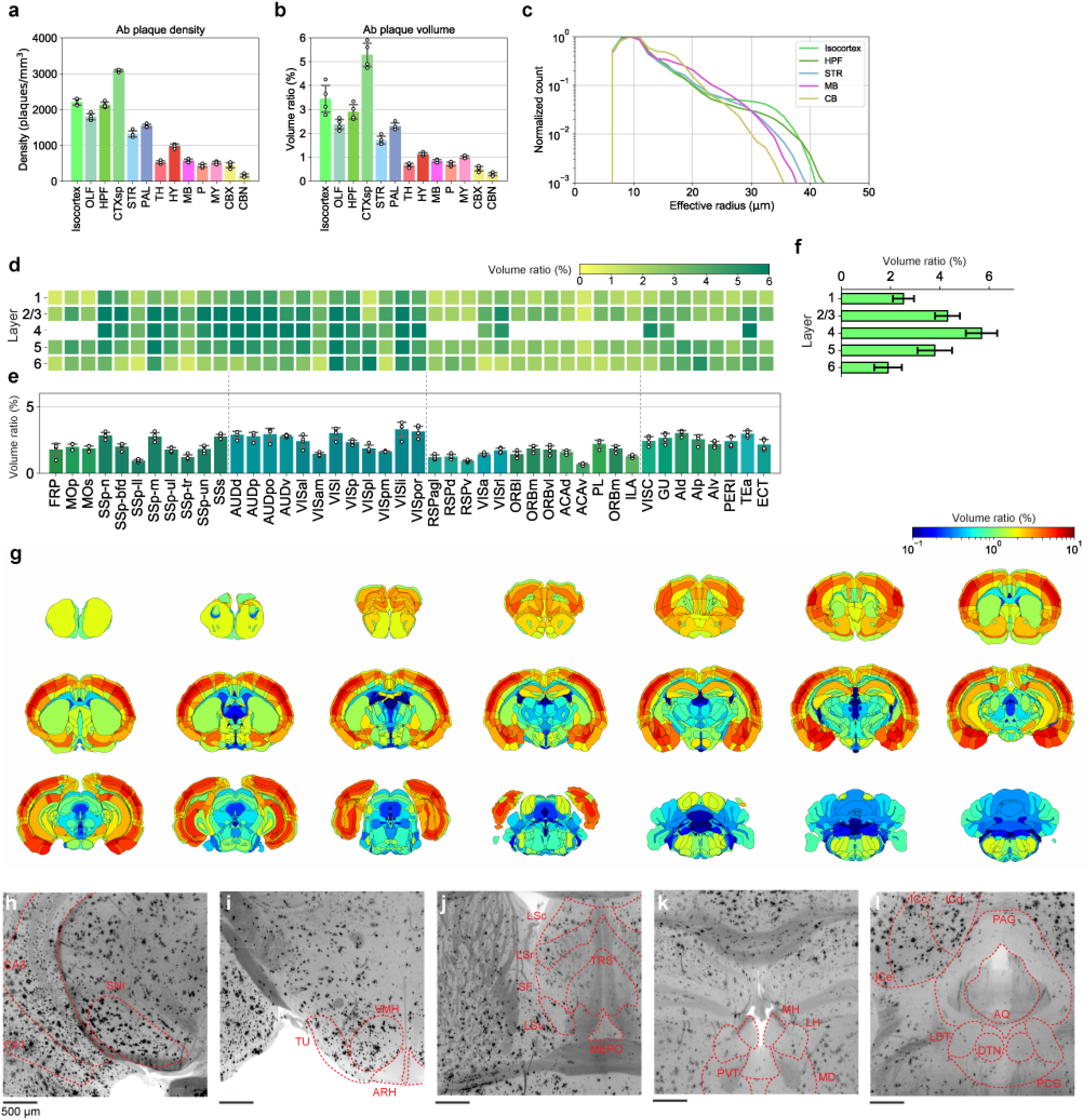
Whole-brain Analysis of Aβ Plaques Accumulation in AD Model Mouse Brain. Using *App*^NL-G-F/NL-G-F^ AD model mouse brain (9-to 10-months-old), brain-wide accumulation of Aβ plaques were quantified by applying wholemount 3D immunostaining and by using CUBIC-Cloud analysis framework. **a**, Density of Aβ plaques (number of plaques/mm^3^) in major brain divisions (*n* = 4). Data are shown as mean ± STD (*n* = 4). **b**, Volume ratio of Aβ plaque in major brain divisions (*n* = 4), computed as (total plaque volume in the region)/(region volume). Data are shown as mean ± STD (*n* = 4). **c**, Distribution of effective radius of Aβ plaques in the isocortex, hippocampus (HPF), striatum (STR), midbrain (MB) and cerebellum (CB) (*n* = 4). **d,e**, The volume ratio of Aβ plaques in the isocortex (*n* = 4). In **e**, data are shown as mean ± STD. **f**, Layer-wise average of **d**. Data are shown as mean ± STD (*n* = 4). **g**, Cartoon heatmap showing the Aβ plaque volume ratio in each brain region (*n* = 4). **h-l**, Raw 6E10 immunostaining images around SNr **h**, VMH **i**, TRS and MEPO **j**, LDT and DTN **k** and MH, LH and PVT **l**.

### Whole-brain analysis of c-Fos expression underling pharmacological sleep induction by LPS

The next important application domain of CUBIC-Cloud is to reconstruct the neuronal activity profile by imaging the protein expressions of immediate early genes (IEGs) such as c-Fos. Such automated analysis would allow comprehensive identification of cellular clusters that underlie an animal’s behavioral phenotype [7, 28, 43, 44]. Reciprocally, one could define an animal’s phenotype in a bottom up manner based on the activity pattern of neuron ensembles. Here, we focused on the effect of LPS. Phenotypically, LPS induces acute sleep in mice, and we thought that IEG-based activity reconstruction would be suitable to track relatively slow dynamics of wake-sleep cycles.

Administration of 150 μg/kg LPS to mice via i.p. injection at CT = 14 caused acute immune response accompanied by prolonged sleep duration, as confirmed by SSS measurement (Supplementary Fig. 5a-c, Methods). In a replicate experiment, brains were sampled 2-3 hours after LPS injection, and the whole-brain c-Fos expression profile was analyzed (Methods). The accuracy of the c-Fos cell detection is shown in Supplementary Fig. 3h. Fig. 4a shows the whole-brain 3D rendering of the detected c-Fos expressing cells (also see Suplementary Table 5). We comprehensively searched for the activated or repressed brain regions by both region-wise and voxel-wise statistical analysis (Supplementary Fig. 5d). Our analysis revealed that c-Fos expression in some of the isocortical areas was reduced (Fig. 4b,c), which included motor and somatosensory areas, presumably reflecting the mouse’s resting state. We also found that some distinct brain nuclei were activated by LPS. Among those, the most notable regions included the BST, PVH, PVT, CEA, PB, NTS and DMX (Fig. 4d).

**Fig. 4.**
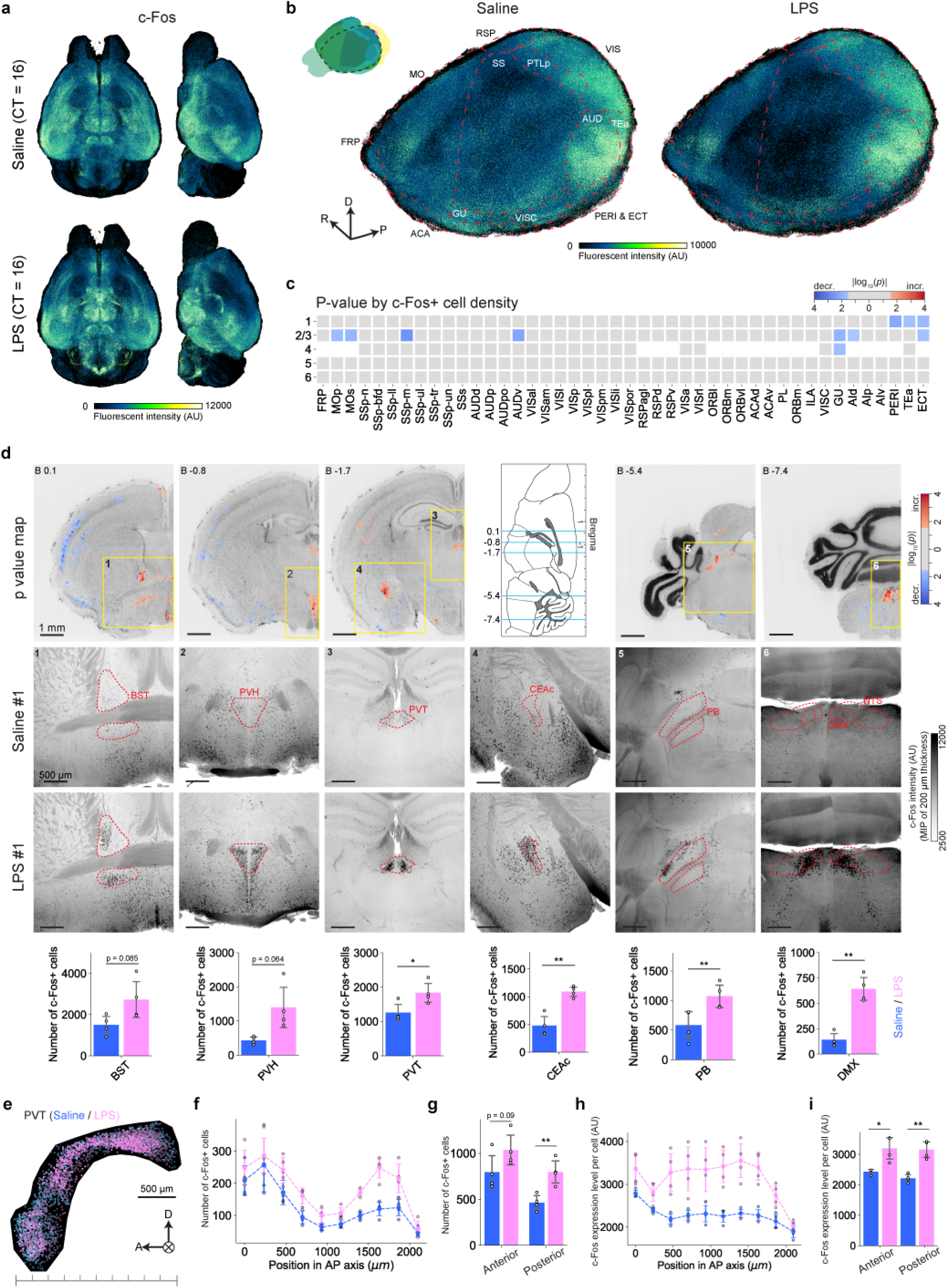
Whole-brain Analysis of c-Fos Expression Level Changes by LPS Administration. LPS acutely induces sleep in mice. Brain-wide neural activity change induced by LPS was quantified by applying whole-mount 3D immunostaining of c-Fos and by using CUBIC-Cloud analysis framework. **a**, Whole-brain views of all cFos+ cells, showing saline (upper) and LPS (lower) administered brains. Each point (i.e. single cell) was assigned a pseudo-color based on its fluorescence intensity. **b**, Magnified 3D view of **a**, where the left isocortex was selectively displayed. Orientation arrows stand for R (right), D (dorsal) and P (posterior). **c**, P-value heatmap showing the isocortex regions whose c-Fos+ cell density was significantly affected by LPS (*n* = 4 each). P-value was computed by comparing the c-Fos+ cell count. The color lookup table is log scaled (base 10), where red color represents the regions that were activated (i.e. more c-Fos+ cells) by MK-801, and blue represents the repressed regions. Regions with no statistical significance (*p* > 0.05) were assigned a gray color. **d**, Distinct brain regions activated by LPS (*n* = 4 each). The top row shows the voxel-wise p-value map. Color lookup table follows that of **c**. The second and third rows are the raw c-Fos images of saline- and LPS-administered group, respectively. The forth row shows the number of c-Fos+ cells of the identified regions. **e**, Plot of c-Fos+ cells in PVT. Cells are pseudo-colored with their intensity values. Pink (blue) dots are from LPS (saline) administered brains, respectively. **f**, The number of c-Fos+ cells in the PVT in 10 divisions along the anterior-posterior (AP) axis. **g**, The number of c-Fos+ cells in the anterior and posterior half of the PVT. The boundary between anterior and posterior region was set at the center of the PVT along AP axis. **h**, The c-Fos expression levels per cell in the PVT 10 divisions along the anterior-posterior (AP) axis. **i**, The c-Fos expression levels per cell in the anterior and posterior half of the PVT. \**p* < 0.05, \*\**p* < 0.01, Welch’s t-test. Brain region acronyms follow the ontology defined by the Allen Brain Atlas.

In our result, the specific part of the BST (the oval region; ovBST) was strongly activated by LPS (Fig. 4d). Indeed, according to a recent study, ovBST is responsible for the inflammation-induced aronexia, and it receives inputs from CEA and PB [45]. Our result was able to successfully identify elevated c-Fos expressions in these spatially separated yet functionally related neurons.

We also observed heterogeneous c-Fos activation in PVT. In terms of the number of c-Fos+ cells, the increase in number was more pronounced in the posterior PVT (pPVT) than the anterior PVT (aPVT) (Fig. 4e-g). In terms of the expression level, both pPVT and aPVT showed similar level of increase (Fig. 4h,i). Recently, Gao et al. [46] identified two classes of distinct neurons in PVT. Type I neurons, densely located in the pPVT, responds to aversive stimuli. On the other hand, type II neurons, dominantly located in the aPVT, become silent upon aversive stimuli. It is also reported that the type II neurons were active during sleep. In the pPVT, our observation aligns with the insight by Gao, where the activated population was likely Type I neurons. In the aPVT, our result might reflect the mixed response of the type II neurons, where aversive inflammatory stimuli and induced sleep were both present. It should also be noted that Type I neurons in the pPVT project to CEA, ILA and ACB. We indeed observed that ILA and ACB were weakly activated (Supplementary Fig. 5d).

### Whole-brain Analysis of Input Cells Projecting to ARH^*Kiss1+*^ Neurons

As the third application domain of CUBIC-Cloud, we show brain-wide connectivity analysis using pseudo-typed Rabies virus (RV). To demonstrate this application, we focused on a population of neurons that secrete kisspeptin (a neuropeptide encoded by *Kiss1* gene) located in the ARH, hereafter termed as ARH^*Kiss1+*^. Those neurons were shown to play an important role in reproduction behavior in mammals by regulating pulsatile release of gonadotrophin-releasing hormone (GnRH) at around 0.3 to 1.0 pulses per hour [47]. Intriguingly, the pulse frequency changes through estrus cycle in females but not in males. As such, we comprehensively investigated the neural inputs to ARH^*Kiss1+*^ neurons to search the mechanism of pulse generation/modulation on the basis of neural circuitry.

To achieve cell-type specific targeting of virus infection, we used the Cre/loxP system and RV trans-synaptic tracing combined with Cre-dependent AAV vectors [48] (Fig. 5a, Methods). After virus injection, brains (both male and female) were cleared by CUBIC reagents and analyzed by CUBIC-Cloud pipeline (Methods). We first checked the distribution of starter cells (GFP+ and mCherry+) to ensure that the injection was successful and the starter cells were well confined within ARH (Fig. 5b). In the present study, the criteria in selecting successful samples was defined as more than 45% of starter cells were localized in ARH or PVp. (Note that area annotated as PVp in the Allen Brain Atlas belongs to a part of ARH in the Paxinos atlas [49].) With this criteria, out of 20 injections, *n* = 3 and *n* = 4 brains were assessed as successful for male and female, respectively (Fig. 5b).

**Fig. 5.**
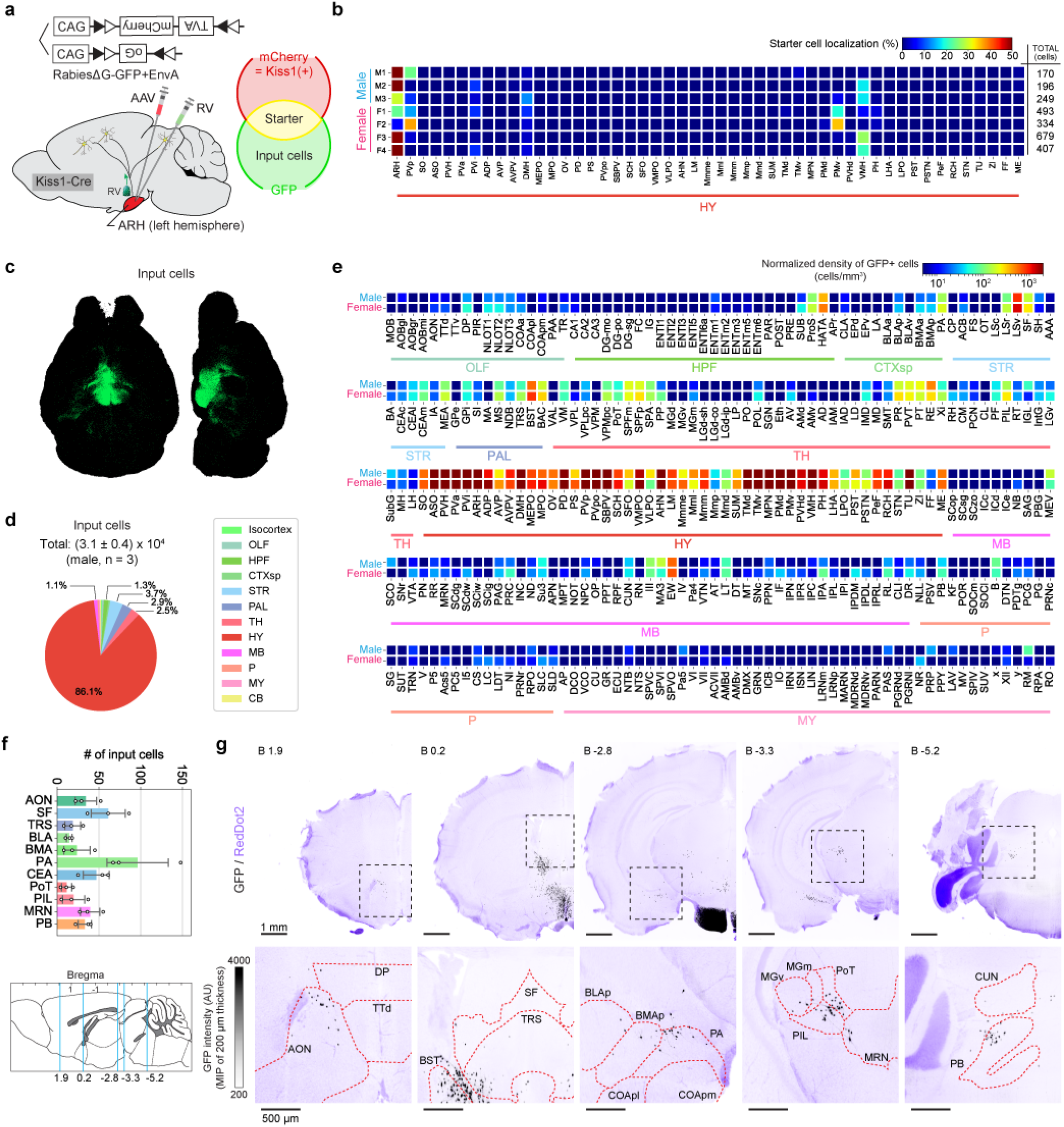
Whole-brain Analysis of Input Cell Populations Projecting to ARH^*Kiss1+*^ Neurons. **a**, Virus injection scheme. AAV carrying mCherry, TVA receptor and optimized glycoprotein (oG) was injected to ARH of Kiss1-Cre transgenic mouse, followed by injection of modified Rabies virus carrying GFP. Cells expressing both mCherry and GFP are the starter cells. **b**, Quantification of starter cell localization. The ratio was computed by dividing the cell count in each region by the total number of starter cells. The total number of starter cells each sample is shown on the right end of the heatmap. **c**, Whole-brain view of all input cells. **d**, Total cell count and the distribution of input cells. Only male brains were considered here. **e**, Cell density heatmap of all brain regions (excluding the isocortex and cerebellum, where virtually no input cells were detected). The mean of male and female brains are shown. **f**, The plot shows extremely sparse input cell populations in previously unidentified brain regions (*n* = 3). Only male brains were considered here. **g**, Raw GFP (black) and nuclear staining (RedDot2, purple) images showing the regions identified in **f**. Macro view (top) and zoomed-in view (bottom) are shown. Brain region acronyms follow the ontology defined by the Allen Brain Atlas.

Fig. 5c shows the whole-brain overview of all input (GFP+ and mCherry-) cells. The accuracy of the GFP+ cell detection is shown in Supplementary Fig. 3b. Our quantitative analysis identified (3.1 ±0.5) × 10^4^ input cells in the male brain (*n* = 3), the majority of which (> 85%) were located within the hypothalamus (Fig. 5d, Suplementary Table 6). As is shown in Fig. 5e, ARH^*Kiss1+*^ neurons receive inputs from dozens of discrete structures throughout the forebrain and brainstem, including the striatum (LS), pallidum (BST), thalamus (PVT), hypothalamus (MPO, MPN, AHN, PVH, DMH, VMH and PH), hippocampal formation (HATA and SUB), midbrain (MRN and PAG), and pons (PB). Remarkably, extremely sparse populations, only a few dozens of cells per region, were reproducibly identified (Fig. 5f,g). In terms of cell density, those populations were often equivalent to less than 10 cell/mm^3^, which could be easily overlooked with slice-based approaches. These sparse populations were not reported in the past literature [50].

We next performed statistical analysis comparing the number of input cells between male and female brains (Methods). Overall, binary connectivity differences (i.e. zero in one sex and some finite number in the other sex) were not observed (Supplementary Fig. 6a). Weak differences were suggested in LS, MPO, MPN and AVP (Supplementary Fig. 6b-h, Methods), which are neighboring with each other. The difference was most pronounced in LSr, which is known to inhibit the lordosis behavior during mating interactions [51]. The sexually dimorphic circuit from LSr to PAG is reported, where female brains contain more neurons in LSr that project to PAG [52]. Our result suggest that LSr sends sexually dimorphic projection to ARH^*Kiss1+*^. The identities of these populations can be fully characterized in the future studies.

## Discussion

In this study, we presented an integrated computational framework for single-cell-resolution whole-brain analysis, named CUBIC-Cloud. Inspired by the data scientific platforms developed in genomics, we postulated that the framework should provide (1) a reference brain atlas and automatic mapping to the reference, (2) open opportunities for the research community to contribute new data, (3) a toolkit to visualize and quantify data, and (4) easy and universal accessibility. As we have shown in this study, CUBIC-Cloud addressed these requirements by designing a new software stack embracing the latest cloud technologies, widely available for researchers in neuroscience (Fig. 1). Users can access CUBIC-Cloud at https://cubic-cioud.com.

After describing the software infrastructures offered by CUBIC-Cloud, we extensively evaluated and validated the accuracy and reproducibility of the proposed framework by various applications. First, we quantified the exact number of various cell types, including PV, SST, ChAT, Th and Iba1 expressing cells (Fig. 2). We were surprised by the small variation between individual animals in terms of labelled cell numbers (quantified SD usually less than 10%), which not only highlighted the accuracy of our analysis but also implied the intricate regulation on cell proliferation in the brain. We further showed that CUBIC-Cloud can also be used to quantitatively understand the pathological state of the AD model mouse (Fig. 3). Together, these demonstrations encourage the future application of CUBIC-Cloud to construct the datasets of the disease models and the drug effects, where decreases or increases of particular markers in specific brain regions may correlate with disease progressions.

We have also explored the application of CUBIC-Cloud to reconstruct the whole-brain neuronal activity profile using c-Fos immunostaining. Our whole-brain analysis showed that administration of LPS, which induces sleepiness, repressed the c-Fos expression in some of the cortical areas, while it activated several distinct brain regions (Fig. 4). For example, our analysis revealed that a distinct subpopulation in the ovBST strongly responded to LPS, and that the response in PVT neurons are heterogeneous. CUBIC-Cloud facilitates to share such knowledge through widely accessible data repository, and users will know exactly, with ≤ 100 *μ*m spacial resolution in 3D space, where the concerning region is. By iterating such experiments, CUBIC-Cloud opens up the possibility to functionally dissect the mouse brain with much finer detail and specificity.

Furthermore, CUBIC-Cloud was used to comprehensively identify brain-wide neuronal connections. As a proof-of-concept demonstration, input of neuronal populations projecting to ARH^*Kiss1+*^ neurons were investigated by using Rabies virus tracers (Fig. 5). Our analysis was successful in identifying extremely sparse populations (less than 10 cells/mm^3^) with high reproducibility. Our result also implied sexually dimorphic projection from LSr to ARH^*Kiss1+*^. Again, we emphasize that CUBIC-Cloud can serve as a central hub for researchers to share such anatomical knowledge.

The above analysis results are all openly accessible at CUBIC-Cloud. The raw image data, along with the markers highlighting the detected cells, are available through CATMAID image browser [53], hosted at http://cubic-atlas.riken.jp.

CUBIC-Cloud is, to our knowledge, the first cloud-based framework to share and publish whole-mouse-brain datasets. We believe, however, that several technical and administrative advancements need to be made to render it a fully functional data scientific platform for neuroscience. Firstly, sharing and viewing of the raw image data has not been addressed in CUBIC-Cloud. There are several software and repository sites for sharing and viewing large biological image datasets [54, 55, 56]. The challenge would be to define an ontology to effectively tag each data so that easy yet flexible search is facilitated. In addition, the deposited image should preferably be aligned with the reference to allow virtual overlay of multiple brain images. To ensure the persistence of the deposited data, involvement of the public agencies may be critical, as is the case in GenBank pr PDB.

Secondly, the current implementation of CUBIC-Cloud asks the user to perform cell detection locally. The cell (object) detection routine, along with the machine learning training, should be integrated in the cloud in the future developments. Indeed, there are already several reports on embracing cloud computing to analyze massive light and electron microscopy images [57, 58, 59, 60]. Interesting bonus of running object detection in the cloud is that the training datasets can also be shared, which will allow to train more general and accurate deep learning models [59]. Furthermore, CUBIC-Cloud is currently aimed at analysing single-cell-resolution image data, where signals of interest are given as dots localized in cell nuclei/soma. Inclusion of object detection routine in the cloud would expand the scope to include fiber and synaptic structures. Technically, such high-resolution image data can be readily obtained by state-of-the-art LSFM, with boosted resolution by expansion microscopy [61, 62, 63, 18]. Future software infrastructure developments, including ones discussed here, will pave the new science towards bottom-up and data-driven elucidation of neuronal functions and circuitry.

**Table 1:** Brain region acronyms used in this paper

| Acronym | Full name |
| --- | --- |
| AAA | Anterior amygdalar area |
| ACB | Nucleus accumbens |
| ADP | Anterodorsal preoptic nucleus |
| AHN | Anterior hypothalamic nucleus |
| ARH | Arcuate hypothalamic nucleus |
| AVP | Anteroventral preoptic nucleus |
| BST | Bed nuclei of the stria terminalis |
| CEA | Central amygdalar nucleus |
| CLI | Central linear nucleus raphe |
| DMH | Dorsomedial nucleus of the hypothalamus |
| DMX | Dorsal motor nucleus of the vagus nerve |
| DTN | Dorsal tegmental nucleus |
| FL | Flocculus |
| HATA | Hippocampo-amygdalar transition area |
| IA | Intercalated amygdalar nucleus |
| IC | Inferior colliculus |
| ILA | Infralimbic area |
| IO | Inferior olivary complex |
| LC | Locus ceruleus |
| LDT | Laterodorsal tegmental nucleus |
| LH | Lateral habenula |
| LRN | Lateral reticular nucleus |
| LS | Lateral septal nucleus |
| LPO | Lateral preoptic area |
| MEA | Medial amygdalar nucleus |
| MH | Medial habenula |
| MRN | Midbrain reticular nucleus |
| MPN | Medial preoptic nucleus |
| MPO | Medial preoptic area |
| NLL | Nucleus of the lateral lemniscus |
| NOD | Nodulus (X) |
| NTB | Nucleus of the trapezoid body |
| NTS | Nucleus of the solitary tract |
| PAG | Periaqueductal gray |
| PB | Parabrachial nucleus |
| PH | Posterior hypothalamic nucleus |
| PP | Peripeduncular nucleus |
| PVH | Paraventricular hypothalamic nucleus |
| PVa | Periventricular hypothalamic nucleus, anterior part |
| PVi | Periventricular hypothalamic nucleus, intermediate part |
| PVp | Periventricular hypothalamic nucleus, posterior part |
| PVpo | Periventricular hypothalamic nucleus, preoptic part |
| PVT | Paraventricular nucleus of the thalamus |
| RAmb | Midbrain raphe nuclei |
| RL | Rostral linear nucleus raphe |
| RR | Midbrain reticular nucleus, retrorubral area |
| RT | Reticular nucleus of the thalamus |
| SFO | Subfornical organ |
| SNc | Substantia nigra, compact part |
| SNr | Substantia nigra, reticular part |
| SO | Supraoptic nucleus |
| SOC | Superior olivary complex |
| SUB | Subiculum |
| TRS | Triangular nucleus of septum |
| VMH | Ventromedial hypothalamic nucleus |
| VTA | Ventral tegmental area |
| ZI | Zona incerta |

## Acknowledgements

Authors would like to thank K. Wilkins and T. Miyawaki for proofreading the manuscript, T.C. Murakami for discussions on CUBIC-Atlas, M. Kuroda for technical assistance on LSFM imaging, S.I. Kubota for helping LPS experiments, R. Tanaka for supporting sample preparations. The App^NL-G-F/NL-G-F^ knock-in mouse strain (RBRC06344) was provided by RIKEN BRC (deposited by T. Saito and T.C. Saido) through the National BioResource Project of the MEXT/AMED, Japan. This work was supported by Grant-in-Aid for JSPS Fellows (JSPS KAKENHI, grant no. 19J13071, to T.M.); ANRI Scholarship (to T.M.); PRESTO (JST, grant no. JPMJPR15F4, to E.A.S.); Grants-in-Aid for Scientific Research (B) (JSPS KAKENHI, grant no. 19H03413 to E.A.S.); Grants-in-Aid for Scientific Research on Innovative Areas (JSPS KAKENHI, grant no. 17H06328 to E.A.S.); ERATO Touhara Chemosensory Signal Project (JST, grant no. JPMJER1202, to K.T.); Grant-in-Aid for Scientific Research (S) (JSPS KAKENHI, grant no. 25221004, to H.R.U.); Brain/MINDS (AMED/MEXT, grant no. JP17DM0207049, to H.R.U.); the Science and Technology Platform Program for Advanced Biological Medicine (AMED/MEXT to H.R.U.); Grants-in-Aid from the Human Frontier Science Program (to H.R.U.); Grants-in-Aid from the Takeda Science Foundation (to E.A.S. and H.R.U.). The graphical user interfaces of CUBIC-Cloud was co-developed by CUBICStars and Tecotec Inc.

## Author contributions

T.M. and H.R.U. designed the study. T.M. created and implemented CUBIC-Cloud software stack. T.M. and K.K. performed data analysis. T.M. and E.A.S. performed LSFM imaging. C.S., K.K. and H.O. performed tissue clearing and immunostaining experiments. K.K. performed sleep measurements using SSS. H.O. assisted in the preparation of AD model mice. S.S. and R.G.Y. assisted in the deployment of the cloud server. E.A.S., K.T., K. Murata and K. Miyamichi prepared Cre-mouse, produced virus and performed injection. T.M., S.S. and H.R.U. wrote the manuscript with the inputs from all co-authors.

## Methods

### Mice

All experimental procedures and housing conditions were approved by the Animal Care and Use Committee of The University of Tokyo. *App*^NL-G-F/NL-G-F^ mice were provided by RIKEN BioResource Research Center (RBRC No. RBRC06344) [42]. Kiss1-Cre mouse was purchased from The Jackson Laboratory (Kiss1-tm1.1(cre/EGFP)Stei/J, stock no. 017701). In all experiments, male mice were used unless otherwise specified.

To sample brains, animals were anesthetized by an overdose of pentobarbital (> 100 mg/kg), then transcardially perfused with 10 mL of PBS (pH 7.4) and 20 mL of 4% paraformaldehyde (PFA). The brains were dissected, post-fixed in 4% PFA overnight at 4 °C and stored in PBS.

### Tissue clearing and whole-brain 3D immunostaining

To clear brain tissues, we used second-generation CUBIC protocols [30, 64]. For delipidation, we used CUBIC-L (10 wt% N-butyldiethanolamine, 10 wt% Triton X-100), and for RI matching, either CUBIC-R+(N) (45 wt% Antipyrine, 30 wt% Nicotinamide, 0.5 vol% N-butyldiethanolamine) or CUBIC-R+(M) (45 wt% Antipyrine, 30 wt% N-methylnicotinamide, 0.5 vol% N-butyldiethanolamine) was used. CUBIC-R+(M) was used for samples labelled by fluorescent proteins (FPs) because of its better FP signal retention, and CUBIC-R+(N) was used otherwise. For all brains, nuclear staining (using either SYTOX-G, BOBO-1 or RedDo2) was applied. For whole-mount 3D immunostaining, we followed CUBIC-HV protocol [31]). The details on antibody staining for each experiment can be found in the corresponding sections in Methods. Before imaging, the sample was embedded in a transparent agarose gel so that it could be rigidly mounted on a microscope stage [18, 64].

### LSFM imaging

A custom-built macro-zoom LSFM (named GEMINI system) was used to image cleared brains (the details of this microscope can also be found in [31]). For illumination, the microscope was equipped with 488, 532, 594 and 642 nm diode or DPSS lasers (SOLE-6, Omicron). The laser sheet was generated by a cylindrical lens and the sheet thickness was adjustable by a mechanical slit. For detection, the microscope was equipped with 0.63X macro-zoom objective lens (MVPLAPO 0.63X, Olympus) and 0.63-6.3X variable zoom optics (MVX-ZB10, Olympus). After passing a suitable fluorescence filter, the fluorescence signal was captured by sCMOS camera (Zyla 5.5, Andor).

To achieve homogeneous light-sheet thickness throughout the field of view, several rectangular image strips with shifted illumination focus were obtained and digitally stitched together, similar to the approach used in TLS-SPIM [65, 66]. The width of the rectangular strip was matched with the Rayleigh range of the illumination light-sheet. In our setup, the sheet thickness was approximately 10 μm and the rectangular strip width was 1500 μm, which required 6 image strips to cover the entire brain.

For RV samples, the image voxel size was (X,Y,Z) = (8.25, 8.25, 9.0) μm with 1.2X intermediate zoom optics. For other experiments, the voxel size was (X,Y,Z) = (6.45, 6.45, 7.0) μm with 1.6X intermediate zoom optics. It should be noted that the effective resolution should take into account the moderate tissue expansion (~1.5X) caused by CUBIC-R treatment.

For each dye/FP, the following laser and fluorescence filter pair was used: Alexa 594 [Ex: 594 nm, Em: 641/75 nm bandpass (FF02-641/75-32, Semrock)], Cy3 [Ex: 532 nm, Em: 585/40 nm bandpass (FF01-585/40-32, Semrock)], SYTOX-G, BOBO-1 and GFP [Ex: 488 nm, Em: 520/40 nm bandpass (FF01-520/44-32, Semrock)], RedDot2 [Ex: 642nm, Em: 708/75 nm bandpass (FF01-708/75-32, Semrock)], mCherry [Ex: 594 nm, Em: 628/32 nm bandpass (FF01-628/32-32, Semrock)].

### Imaging the fluorescent bead embedded in cleared tissue

1.0 μm-diameter green-yellow fluorescent beads (Thermo Fisher, #F8765) were diluted in PBS so that the final bead concentration was 0.9 × 10^7^ particles/ml. This bead-mixed PBS solution was perfused in mice, prior to the PFA perfusion. Because the bead surface was modified with amine, PFA was able to cross-link and fix the beads within the tissue. After tissue clearing, the whole brain was imaged using the macro-zoom LSFM with XYZ voxel resolution of 6.45×6.45×7.0 μm (Supplementary Fig. 2a). Then, single and well-isolated bead particles were manually annotated (*n* > 15 for each brain region) using ITK-SNAP software [67]. Subsequently, the mean spot profiles were computed and fitted with Gaussian using custom Python code. The fitted sigma values from six regions were all within 4.4 to 4.9 μm (lateral) and 5.3 to 6.7 μm (axial) (Supplementary Fig. 2b), validating homogeneous image quality throughout the entire brain. Given the digital sampling frequency (6.5 μm) of the microscope used, this result was nearly the ideal PSF.

### Cell detect/on

The cell detection workflow used in this study is shown in Fig. 1a (“step 3”). As the first step, a machine-learning algorithm was used to classify voxels into labelled cells and other structures. For this step, we used ilastik [32]. To train the classifier, for each label type (such as c-Fos), manual annotation images were prepared. In this study, three classes were defined, which were (1) signals of interest, i.e., cells labelled by FPs or antibodies (2) bright but false signals, such as non-specific binding of antibodies to vascular structures or neurites extending from cell bodies and (3) background (i.e. void space). Typically, 5,000 to 10,000 voxels were annotated as class 1 per one dataset. To increase the robustness, at least two brains with identical labelling conditions were annotated. Image annotation was performed using ITK-SNAP software [67].

Next, following the ilastik workflow, image feature descriptors were selected to distinguish the three classes defined above. For PV, Sst, ChAT, Th, Iba1, c-Fos, RV-GFP and AAV-mCherry images, the selected descriptors were Gaussian (σ = 0.3, 0.7 voxel), Gaussian gradient magnitude (σ = 0.7 voxel), difference of Gaussian (σ = 0.7, 1.0, 1.6, 3.5 voxel) and Hessian of Gaussian eigenvalues (σ = 1.0, 1.6, 3.5 voxel). For Aβ images, difference of Gaussian (σ = 5.0 voxel) and Hessian of Gaussian eigenvalues (σ = 5.0 voxel) were additionally included, so that the larger spatial context was taken into account. Then, ilastik software trained the voxel classifier using random forest algorithm. Hyperparameters related to random forest algorithm was automatically configured by ilastik. After the classifier was trained, it was applied to the test dataset to evaluate accuracy. If obvious error was present, manual annotation data was further supplemented, so that the classifier became more robust against such errors. Once the classifier achieves satisfactory accuracy, it could reliably be applied to other images with the same label type.

By applying the voxel classifier trained above to each brain image, a probability image was produced, where the value of each voxel represents the probability of that voxel being class 1 (Supplementary Fig. 3a). The probability value was given in the range [0, 1]. Using this probability image, a custom Python program isolated individual cells in the following way. First, the probability threshold, *P*_th_ = 0.7, was applied to make a binarized image. Then, connected voxels were searched and merged together, to find individual objects. If the identified object volume was larger than a threshold, *V*_th_, it was sent to the object separation routine. The object separation routine simply finds local maxima with an exclusion distance 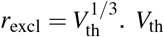 was heuristically determined to be *V*_th_ = 4^3^ = 64 for PV, Sst, c-Fos, Iba1, Rabeis-GFP and AAV-mCherry, and *V*_th_ = 5^3^ = 125 for ChAT and Th. For Aβ plaque segmentation, *V*_th_ = ∞ was used. Hence, there are two free parameters (*P*_th_ and *V*_th_), both of which can be intuitively determined.

Using 20 CPU cores, cell detection of one 3D image stack (typically 2560×2160×1200 voxels) took about 1 hour to complete.

The Python source code and its documentation is available at https://github.com/DSPsleeporg/ecc.

### Accuracy evaluation of cell detection

Accuracy of the above explained cell detection procedure was extensively evaluated by comparing automated counting results with manual cell counting (Supplementary Fig. 3b-h). Manual cell counting was performed by cropping a small cubic image volume (50 or 75 or 100 voxels, depending on the cell density) from brain images which were not used in machine learning training. Well-trained human annotators (*n* = 2) independently marked all of the cells present in the image and typically yielded 100-200 marked cells. Image annotation was performed using ITK-SNAP software [67]. Cells annotated by both human and algorithm were regarded as true positives. Cells annotated by human but not by algorithm was regarded as false negative. Cells annotated by algorithm but not by human were regarded as false positives. Then, true positive rate (TPR, also called sensitivity) and positive predictive value (PPV, also called precision) were evaluated for each human annotator. To quantify the overall performance, F1 score, defined as *F*1 = 2* (*PPV* × *TPR*)/(*PPV* + *TPR*), was also evaluated. For most of the label types and brain regions, our cell detection algorithm robustly demonstrated good F1 scores, with average score being 0.80 (PV), 0.83 (Sst), 0.88 (ChAT), 0.80 (Th), 0.88 (Iba1), 0.83 (c-Fos) and 0.89 (GFP).

Raw 3D image data, along with markers highlighting all detected cells, are available at http://cubic-atlas.riken.jp through an interactive image viewer powered by CATMAID [53].

### CUBIC-Atlas

CUBIC-Cloud uses CUBIC-Atlas version 1.1 [18] as the reference brain coordinate, to which all individual brain data were mapped. From the originally published atlas (version 1.0), we added a few minor updates in this study to generate CUBIC-Atlas version 1.1. The updates included (1) slight discontinuity between dorsal and ventral image volume (so-called “theta tile displacement” in Murakami et al. paper) were corrected and (2) region annotation was updated to be compatible with the Allen Brain Atlas CCFv3 (October 2017). CUBIC-Atlas v1.1 can be downloaded from http://cubic-atlas.riken.jp.

### Brain registration

CUBIC-Cloud uses the symmetric image normalization (SyN) algorithm implemented in ANTs library [34] to run registration between CUBIC-Atlas (“fixed” brain) and individual brain sample (“moving” brain). First, nuclear staining image of the “moving” image was downscaled to a voxel size of 50×50×50 μm. Nuclear staining image of CUBIC-Atlas was downscaled to a voxel size of 80×80×80 μm. Considering the sample’s physical expansion by clearing treatment (2.2X for CUBIC-Atlas and 1.5X for CUBIC-R+ treated brains), this downscaling operation resulted in an effective voxel size of about 35×35×35 μm in both images. The registration first computed affine transformation to coarsely align the orientation and size, using mutual information as the optimizer metric. Subsequently, non-linear warping was computed by SyN algorithm, which optimized the warp field by maximizing the normalized cross-correlation (NCC) between the two images under diffeomorphic regularization [34]. Given image *I*(**x**) and image *J*(**x**), the NCC value between *I* and *J* at the voxel position **x** is given by

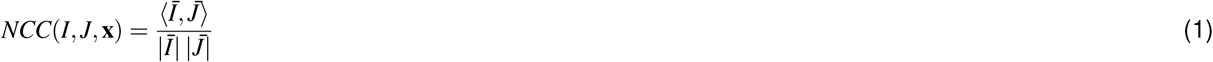

where 〈*A,B*〉 represents the inner product taken over a local window with radius *R* centered at position **x**. |*A*| is the L2 norm of the vector computed over a local window with radius *R*. Here, *I*(**x**) = *I*(**x**) – *μ_I_*(**x**) means the subtraction of the local mean, where local mean *μ_I_*(**x**) is computed over a local window with radius R centered at position **x**. *R* = 4 (voxels) was used throughout our brain registration.

The representative registration result is visualized in Supplementary Fig. 3i, along with the NCC value heatmap (Supplementary Fig. 3l). To show the reproducibility of the registration, 20 individual brains were mapped onto CUBIC-Atlas with identical registration parameters. The mean NCC value of each coronal planes were computed and plotted (Supplementary Fig. 3h). 20 independent curves overlap with each other, meaning that the optimization attempt by the registration reached saturation. NCC value tends to show higher value in the olfactory area and cerebellum, due to the presence of more distinct structural features.

### Architecture of CUBIC-Cloud

CUBIC-Cloud’s entire application stack is deployed on the cloud computing infrastructures offered by Amazon Web Service (AWS). The cloud is constructed using the serverless architecture [68]. Serverless architecture have zero real instances that are always running; instead, the cloud is composed by connecting microservices, which are dynamically invoked by events. Such cloud design eliminates the cost of idling servers, while allowing to flexibly and instantly scale out the computing power when the traffic or load to the service increases.

The schematic illustration of the cloud architecture is shown in Supplementary Fig. 1. When user accesses the web site, the trafic is first handled by CloudFront. CloudFront is responsible for caching static contents, managing SSL/TLS and web application firewall (WAF). Then, static web contents are fetched from S3 bucket and returned to the user. User authentication is handled by Cognito. Once authenticated, users can access the protected API endpoints securely using json web token (JWT). All REST API requests are routed by API Gateway and handled by Lambda. Lambda handlers have access to various back-end resources, including the databases and the data buckets. The metadata of the users, brains, notebooks, and studios are stored in DynamoDB. Large data files (such as images and csv tables) are stored in S3.

Once a user uploads the brain data, the upload completion event from S3 triggers a “preprocessing” task in the ECS cluster. Preprocessing includes brain registration, transformation, and data conversion. ECS automatically launches a new EC2 instance, pulls the Docker container from ECR registry, and initiate a new task. The task execution is orchestrated by StepFunctions. Notebook tasks (i.e. generating plots) are similarly orchestrated by StepFunctions, except that the runtime is either Lambda or Fargate, depending on the required memory size of the task.

Most of the API handlers are written in Python. The cloud resources are managed by AWS CDK library for Python, and the cloud is deployed using the CloudFormation generated by the CDK library. The user interfaces (UIs) were constructed using HTML/CSS/JavaScript and Vue.js framework.

### A web-based whole-brain viewer

CUBIC-Cloud offers a point-cloud based interactive 3D brain viewer, a feature called studio. The viewer is written in JavaScript, and runs in most of the standard web browsers, including Google Chrome and Firefox. It uses WebGL for hardware accelerated 3D rendering. The core of the point cloud rendering engine was adopted from the open-source project, Potree [69]. Following Potree, CloudEye uses a specialized point cloud format where the whole point cloud was divided and stored in a multi-resolution hierarchical structure (octree structure). This hierarchical data structure enabled the adaptive data querying in response to client’s viewpoint, in which a portion of the points near the viewer’s camera was selectively loaded. The octree-formatted point cloud data was automatically generated as a part of the preprocessing task in the CUBIC-Cloud server. Each point can be attached with several attributes, including the region ID and fluorescent intensity. Points may be colored using these attributes. For example, points may be given gradient colors based on their fluorescence intensity values.

Users can navigate and explore the whole-brain data by intuitive mouse interactions. Points can be selectively hidden/displayed based on the region ID. Arbitrary combinations of brains may be overlaid with user-defined colors. Users can also make a slice view, which can be moved or rotated with simple mouse dragging. Users can also grab a point (cell) and query its information, such as fluorescent intensity.

### Whole-brain analysis of PV and SST expressing neurons

C57BL/6N wild-type mice brains (8-week-old, *n* = 4) were cleared, stained, imaged and analyzed as described in the corresponding sections in Methods. Brains were stained with PV antibody (Swant, #PV235; 1/50 dilution; anti-mouse IgG1 secondary Fab fragment conjugated with Cy3 (Jackson ImmunoResearch, #115-167-185)), SST antibody (Millipore, #MAB354; 1/10 dilution; anti-rat IgG secondary Fab fragment conjugated with Alexa 594 (Jackson ImmunoResearch, #112-587-008)) and nuclear staining dye (BOBO-1, Thermo Fisher #B3582). The whole-brain summary of detected PV+ and SST+ cells are provided in Suplementary Table 2.

We discussed the PV+ and SST+ cell in the isocortex in the main text. Within the striatum, PV+ cells were almost entirely absent in the LS and anterior, central, intercalated and medial amygdalar nucleus (AAA, CEA, IA and MEA), as observed previously [70]. Distribution in the brain stem faithfully reproduced the previous slice-based immunohistochemical studies [71, 72] (Fig. 2l). In general, the thalamus contained low numbers of PV+ or SST+ cells, except that dense PV+ cells were present in the RT and PP. In the hypothalamus, although PV+ cells were sparse, many nuclei contained medium to high density of SST+ neurons. Within the midbrain, PV+ cells were particularly abundant in the IC and SNr, while SST+ cells were most frequently observed in the RAmb. Within the pons and medulla, the NTB, SOC and NLL contained both PV+ and SST+ cells with relatively high density, while sparsely scattered populations were observed in other areas. In the cerebellum, there were a large number of PV+ neurons in Purkinje layers. Distinct SST+ cell clusters were found in the NOD and FL.

### Whole-brain analysis of ChAT expressing neurons

C57BL/6N wild-type mice brains (8-week-old, *n* = 4) were cleared, stained, imaged and analyzed as described in the corresponding sections in Methods. Brains were stained with ChAT antibody (abcam, #ab178850; 1/200 dilution; anti-rabbit IgG secondary Fab fragment conjugated with Alexa 594 (Jackson ImmunoResearch, #111-587-008)) and nuclear staining (SYTOX-G, Thermo Fisher, #S7020). The whole-brain summary of detected ChAT+ cells are provided in Suplementary Table 2.

About half of ChAT+ cells were concentrated in striatum and pallidum (34.5% and 13.8%, respectively). Continuously spreading from these regions, some ChAT+ cells were present in the hypothalamus, including lateral, medial and anteroventral preoptic areas (LPO, MPO, AVP) and SO. ChAT+ neurons were hardly observed in the olfactory area, hippocampus, cortical subplate and thalamus. These observations were in good agreement with the slice-based immunohistochemical study by [73].

### Whole-brain analysis of Th expressing neurons

C57BL/6N wild-type mice brains (8-week-old, *n* = 4) were cleared, stained, imaged and analyzed as described in the corresponding sections in Methods. Brains were stained with Th antibody (Santa Cruz Biotechnology, #sc-25269; 1/20 dilution; anti-mouse IgG2a secondary Fab fragment conjugated with Alexa 594 (Jackson ImmunoResearch, #115-587-186)) and nuclear staining (SYTOX-G). The whole-brain summary of detected Th+ cells are provided in Suplementary Table 2.

The majority of the detected Th neurons were localized in well-known dopaminergic cell groups (A8 to A16) and noradrenergic cell groups (A1 to A7) [74]. Dopaminergic cell groups include the RR, SNc, rostral and central linear nucleus raphe (RL and CLI) and VTA, which form the A8, A9 and A10 in midbrain. Within the hypothalamus, Th neurons were clustered in the periventricular hypothalamic nucleus, anterior, posterior, intermediate and propotic parts (PVa, PVp, PVi, PVpo), ARH, ZI and ADP, which form A11-A15 cell groups. Th neurons were numerous in the olfactory area (A16), selectively localized in the glomerular layer. Noradrenergic cell groups formed distinct bands crossing several nuclei in the medulla and pons, which included the LRN, NTS and DMX, which form A1 and A2. In the pons, a particularly high density was observed in and around LC, which forms A6. No significant population of Th+ cells were observed in the isocortex, hippocampus, cortical subplate, striatum and pallidum.

### Brain nuclei segmentation using alpha shape algorithm

A densely aggregated ChAT expressing cells were found in the LDT and its neighboring regions. To segment this brain region automatically, a rectangular region containing these cells were manually cropped. Then, to remove the isolated cells that was not part of the continuous body of the nuclei, the following filter was applied: for each cell in the ensemble, the number of neighboring cells within the radius 100 μm were counted, and if the count is less than 2, the cell was removed from the ensemble. After this filtering, the polygonal surface enclosing the points was constructed using alpha shape algorithm with alpha radius parameter 300 μm, implemented in MATLAB. This method was applied to *n* = 4 brains aligned with CUBIC-Atlas, obtaining four independent boundary surfaces. Using the same method, cluster of TH expressing cells around LC were segmented (*n* = 4).

Lastly, the overlap between ChAT-defined boundaries were evaluated by computing the Dice’s coefficient, *c* = (2* *V*_*V*1∩*V*_2__)/(*V*_1_ + *V*_2_) where *V*_*V*1∩*V*_2__ is the volume of the overlap between the two polygons whose volumes are *V*_1_ and *V*_2_, respectively. The Dice’s coefficient of all possible pairs, 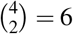, were evaluated. The same analysis was applied to TH-defined boundaries. Lastly, the overlaps between ChAT and TH boundaries were evaluated by picking all possible pairs, 4^2^ = 16.

### Whole-brain analysis of Iba1 expressing cells

C57BL/6N wild-type mice (8-week-old, *n* = 7 for LPS-administered and control group, respectively) were administered with either 1 mg/kg of LPS or saline via i.p. injection. 24 hours after injection, brains were sampled. Subsequently, brains were cleared, stained, imaged and analyzed as described in the corresponding sections in Methods. Brains were stained with Iba1 antibody (Wako, #013-26471; 1/50 dilution; directly conjugated with red dye) and nuclear staining (SYTOX-G). The whole-brain summary of detected Iba1+ cells are provided in Suplementary Table 3.

### Whole-brain analysis of Aβ plaques

*App*^NL-G-F/NL-G-F^ mice (9-to 10-month-old, *n* = 4) were cleared, stained, imaged and analyzed as described in the corresponding sections in Methods. Brains were stained with β-Amyloid (6E10) antibody (Biolegend, #93049; 1/100 dilution; anti-mouse IgG1 secondary Fab fragment conjugated with Alexa 594 (Jackson ImmunoResearch, #115-587-185)) and nuclear staining (SYTOX-G).

### Whole-brain analysis of c-Fos profile change induced by LPS

To monitor the sleep duration of mice in an non-invasive manner, we used a respiration-based sleep staging method, SSS [75]. C57BL/6N wild-type mice (8-week-old, *n* = 4 each for LPS-administered and control group) were housed in a SSS chamber for 3 days (basal measurement; LD cycle) prior to the injection. On the fouth day at ZT = 14, mice were administered 150 μg/kg LPS from Escherichia coli(Sigma-Aldrich, #L2990) via i.p. injection, while control mice were administered saline (Supplementary Fig. 5d). After injection, mice were housed in the SSS chamber for another 24 hours to monitor sleep.

Brains for whole-brain imaging were collected in a replicate experiment with the identical conditions as above, except that mice were housed in DD cycle and that brains were sampled at CT = 16-17 after i.p. injection (*n* = 4 in total for each group, obtained in 1 batch). Collected samples were cleared, stained, imaged and analyzed as described in corresponding sections in Methods. Brains were stained by c-Fos antibody (CST, #2250S; 1/75 dilution; anti-rabbit IgG secondary Fab fragment conjugated with Alexa 594 (Jackson ImmunoResearch, #111-587-008)) and nuclear staining dye (SYTOX-G).

### Rabies virus injection

The following AAV vectors were generated de novo by PENN vector core using the corresponding plasmids. AAV serotype 9 CAG-FLEx-TCb (1.5 × 10^13^ gp/ml) was made using the plasmid described previously [48]. Here TCb stands for TVA-mCherry expression cassette optimized to increase mCherry brightness. To generate AAV serotype 9 CAG-FLEx-oG (4.5 × 10^13^ gp/ml), engineered and optimized glycoprotein (oG) [76] sequence was ligated to pAAV-FLEX sequence from pAAV-FLEX-GFP (Addgene).

Preparation of rabies virus was conducted by using the RVΔG-GFP, B7GG and BHK-EnvA cells as previously described [77]. The EnvA-pseudotyped RVΔG-GFP+EnvA titer was estimated to be 1.0 × 10^9^ infectious particles/ml based on serial dilutions of the virus stock followed by infection of the HEK293-TVA800 cell line (a gift from Dr. Edward Callaway at Salk Institute).

For trans-synaptic tracing using rabies virus, about 20 nL of mixture of AAV9 CAG-FLEx-TCb and CAG-FLEx-oG (diluted to 1.5 × 10^12^ gp/ml each) was injected into the ARH of Kiss1-Cre mice. The first AAV transduced a TVA receptor (fused with mCherry) for EnvA. The second AAV transduced RV glycoprotein (oG) playing a predominant role in the trans-synaptic transport of RV. The injection coordinate was P1.1, L0.2, V5.9 (distance in mm from the Bregma for the posterior [P], and lateral left [L] positions and from the brain surface for the ventral [V] position). Three weeks later, 30 nL of Rabies ΔG-GFP+EnvA was injected into the same brain region to initiate trans-synaptic tracing. Because there is no cognate receptor for EnvA in the mouse brain, RVΔG+EnvA only infects TVA-expressing cells. oG expression from the second AAV complements the RVΔG, allowing retrograde monosynaptic tracing from Cre-expressing cells. Seven days later, brains were sampled for CUBIC treatment.

### Whole-brain analysis of RV-injected brains

After AAV and RV injection, Kiss1-Cre mice (13- or 14-week-old at the time of brain sampling) were cleared, stained, imaged and analyzed as described in the corresponding sections in Methods. Brains were stained with nuclear staining (RedDot2, Biotium, #40061).

As a negative control experiment, AAV and RV injection was performed using BALB/c wild-type mice brain (*n* = 3). After clearing, the whole-brain image was obtained by LSFM. No GFP or mCherry signals were observed by manual inspection, confirming the absence of Cre-indepentent leakage of AAV vectors and specificity of the virus delivery.

Starter cells were searched by identifying dual positive (mCherry+ and GFP+) cells. For each channel, cell counting was independently performed, and the center of the mass of the detected cell was obtained. For each mCherry+ cells, if a GFP+ cell was present within a distance of 24 μm, the cell was counted as starter. Note that because the cleared tissue was expanded by a factor of ~1.5, this was roughly 16 μm in untreated tissue. Occasionally, slight voxel shift (typically no more than 4 voxels) occurred between GFP and mCherry channels, which was presumably caused by slight misalignment between the 488 nm and 594 nm illumination laser or re-focusing of the microscope. To correct this, small 3D volumes with distinct features (typically (X,Y,Z) = (50,50,20) voxel volume, *n* = 4 or *n* = 3) from mCherry and GFP channels were cropped, and the voxel shift was computed by registering two images using ANTs, where transformation was restricted to only translation. Then, the cell coordinates were corrected by the mean of the computed shift.

To carry out statistical analysis of input cell numbers between male and female brains, we used the normalized cell count, *n*_norm,*i*_, where *i* represents the ID of the brain region. If we let the raw cell number of each brain region be *n*_raw,*i*_, *n*_norm,*i*_ is simply expressed as *n*_norm,*i*_ = *n*_raw,*i*_/(∑_*i*_ *n*_raw,*i*_).

## Data availability

Raw 3D image data analyzed in this study are available at http://cubic-atlas.riken.jp through an interactive image viewer powered by CATMAID [53]. The analyzed whole-brain data in point cloud format is deposited on http://cubic-atlas.riken.jp, as well as CUBIC-Cloud’s public repository. The additional data that support the findings of this study are available from the corresponding author upon reasonable request.

## Code availability

Cell detection program used in this study is available at GitHub code repository (https://github.com/DSPsleeporg/ecc). CUBIC-Cloud computing service is accessible at https://cubic-cloud.com.

CUBIC-Cloud is free for use for viewing data. Requests to upload brains or create notebooks are handled for a fee, to cover and compensate the maintenance cost of the cloud server.

---

**Supplementary Figure 1.**
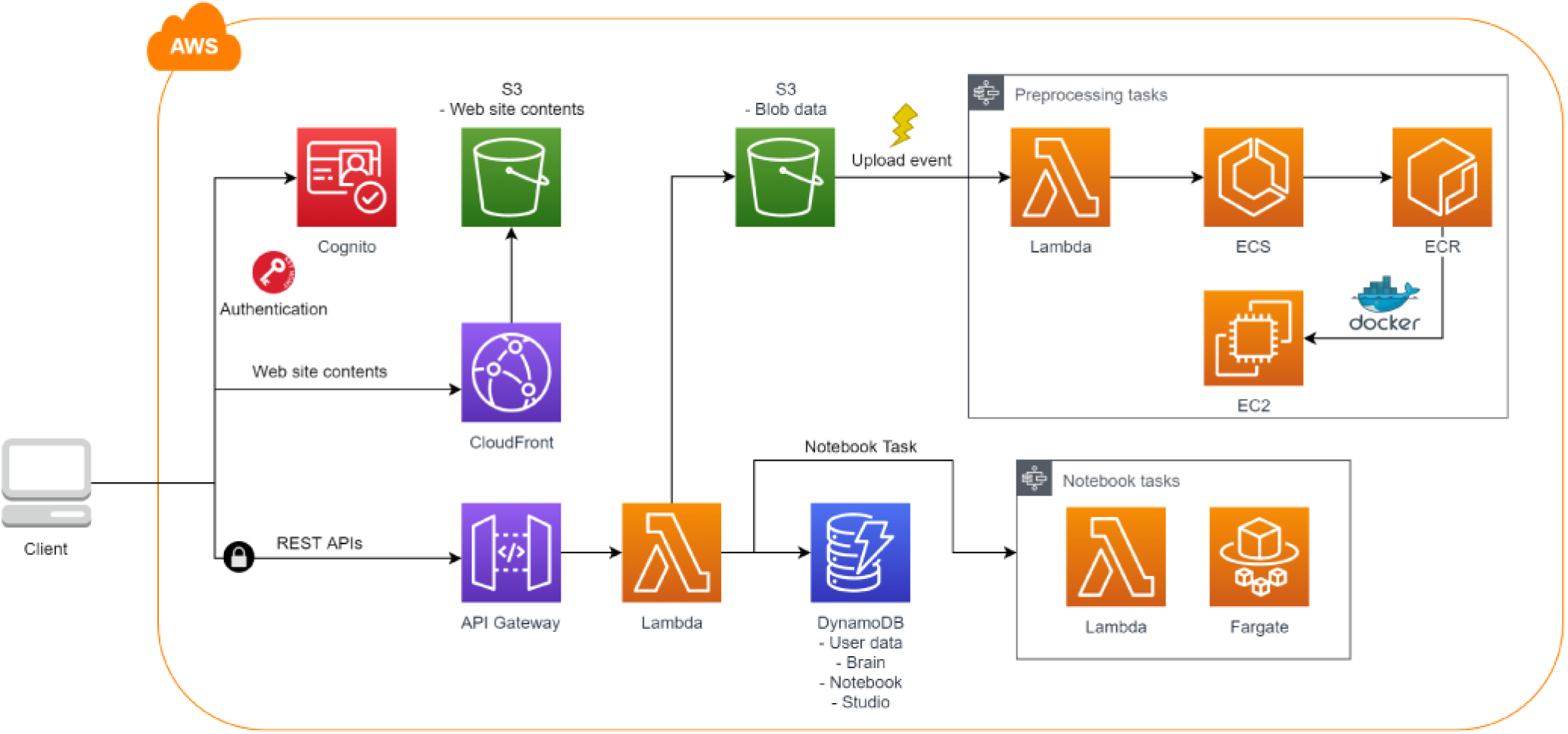
Architecture of CUBIC-Cloud. Schematic diagram showing the architecture of CUBIC-Cloud (see Methods).

**Supplementary Figure 2.**
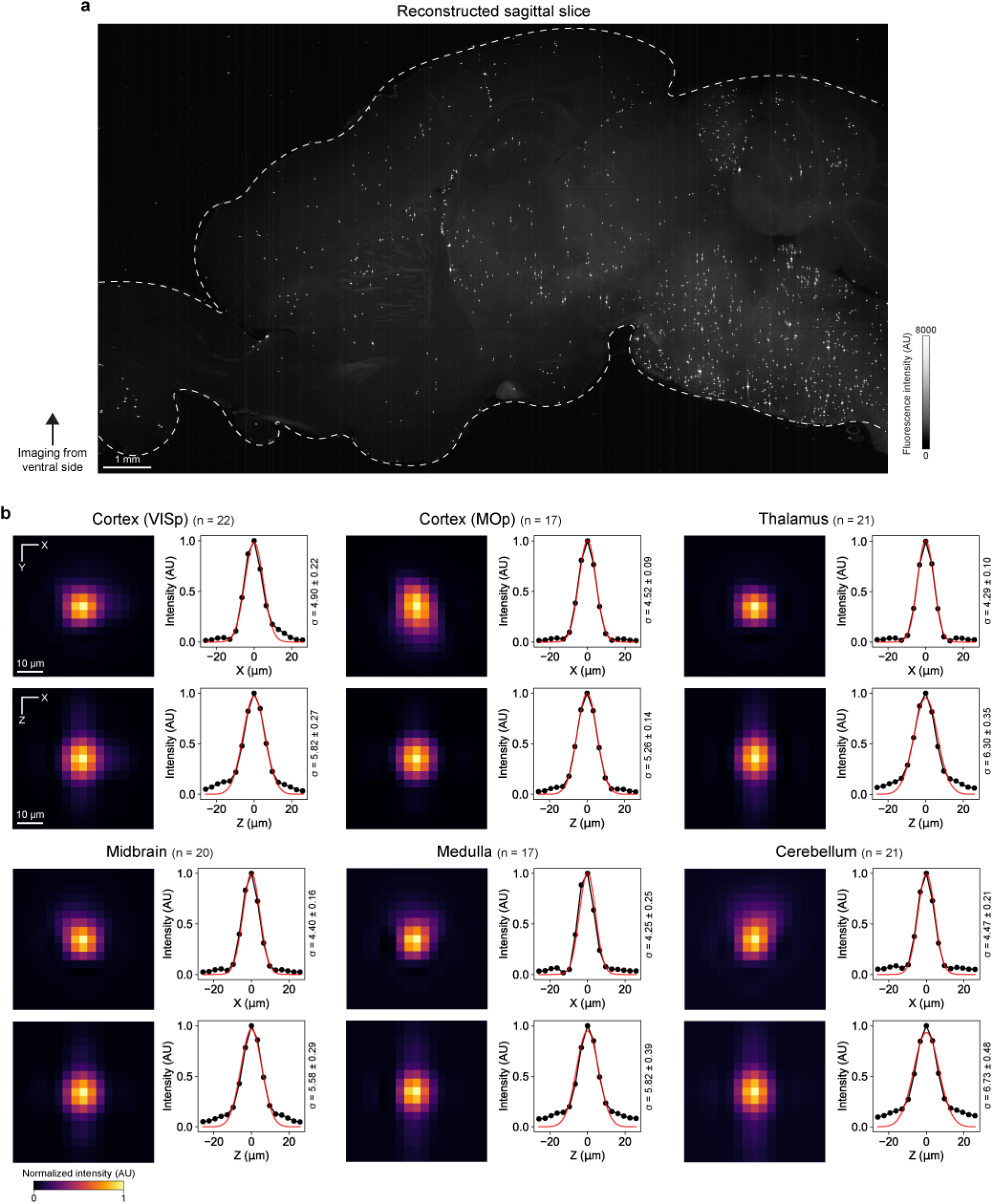
Imaging 1.0 μm-diameter Fluorescent Beads Embedded in Cleared Tissue. 1.0 μm-diameter fluorescent beads embedded in cleared tissue were imaged using macro-zoom LSFM (1X magnification, 6.5 μm voxel size) from the ventral side (see Methods). **a**, Representative sagittal slice (virtually reconstructed from horizontal-major 3D image stack) showing the fluorescent beads embedded in tissue. **b**, Bead spot profiles measured in six brain regions. Lateral and axial profiles were fitted with Gaussian, respectively, and the fitted curves are shown along with the raw data points (see Methods). The fitted sigma values (with 95% confidence interval) of the Gaussian are also shown. The number of particles used to average are shown in the graph. Brain region acronyms follow the ontology defined by Allen Brain Atlas.

**Supplementary Figure 3.**
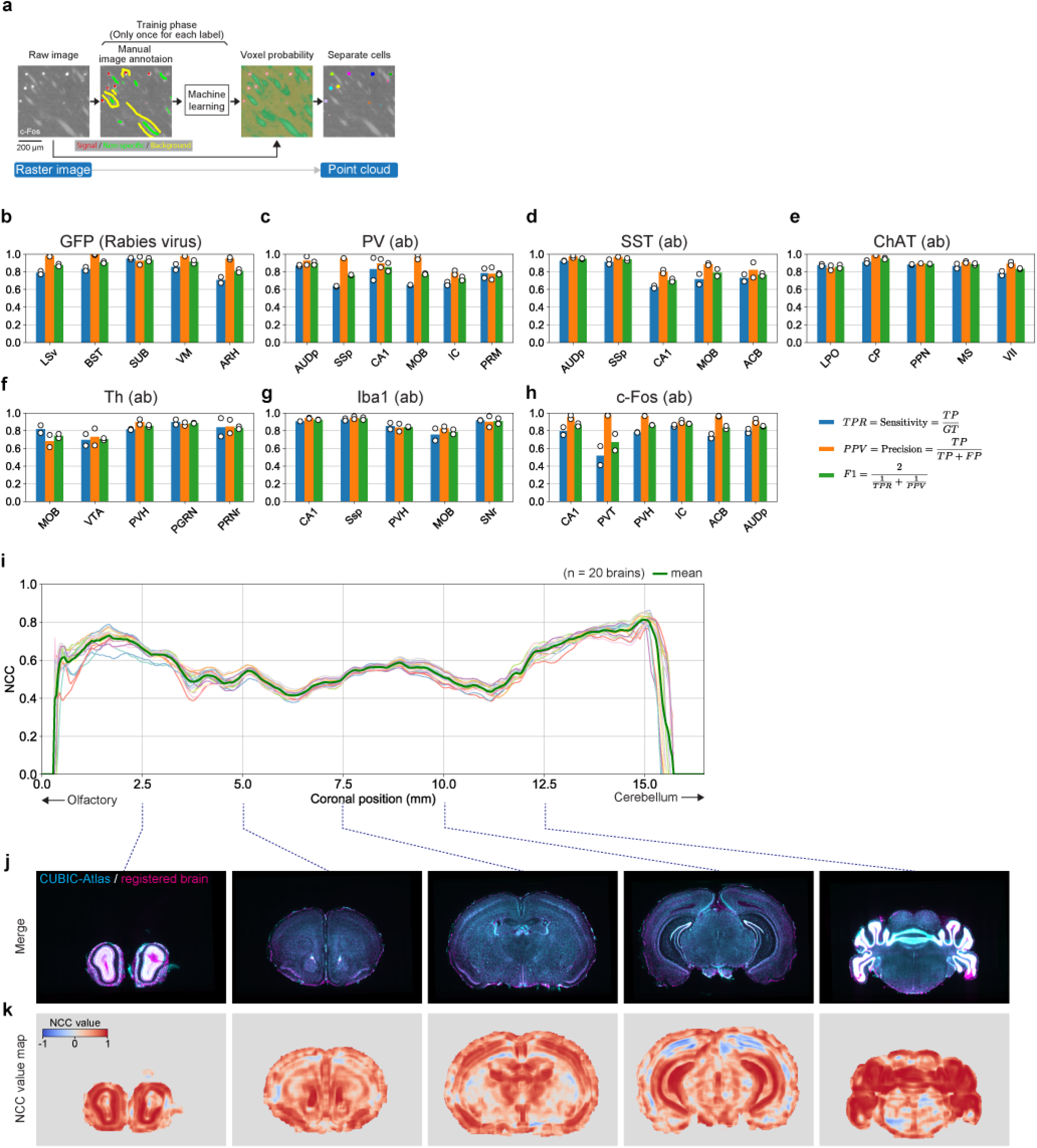
Validation of Cell Counting and Registration Methods. **a**, The cell detection workflow overview (see Methods). **b-h**, Accuracy evaluation of the cell counting algorithm. True positive rate (TPR), positive predictive value (PPV) and F1 score were evaluated for seven label types in more than five brain regions. Ground truth was prepared by two independent annotators (see Methods). **i**, Normalized cross-correlation (NCC) value between two brains after registration. Mean NCC value of each coronal slice are plotted. 20 brains from different mice were independently registered onto CUBIC-Atlas. Individual profiles (thin lines with light colors) as well as the mean (thick green line) are shown. **j**, Representative brain registration result. CUBIC-Atlas (cyan) and registered brain (magenta) are overlaid. **k**, Voxel-wise NCC value map computed for the images shown in **j**. Brain region acronyms follow the ontology defined by Allen Brain Atlas.

**Supplementary Figure 4.**
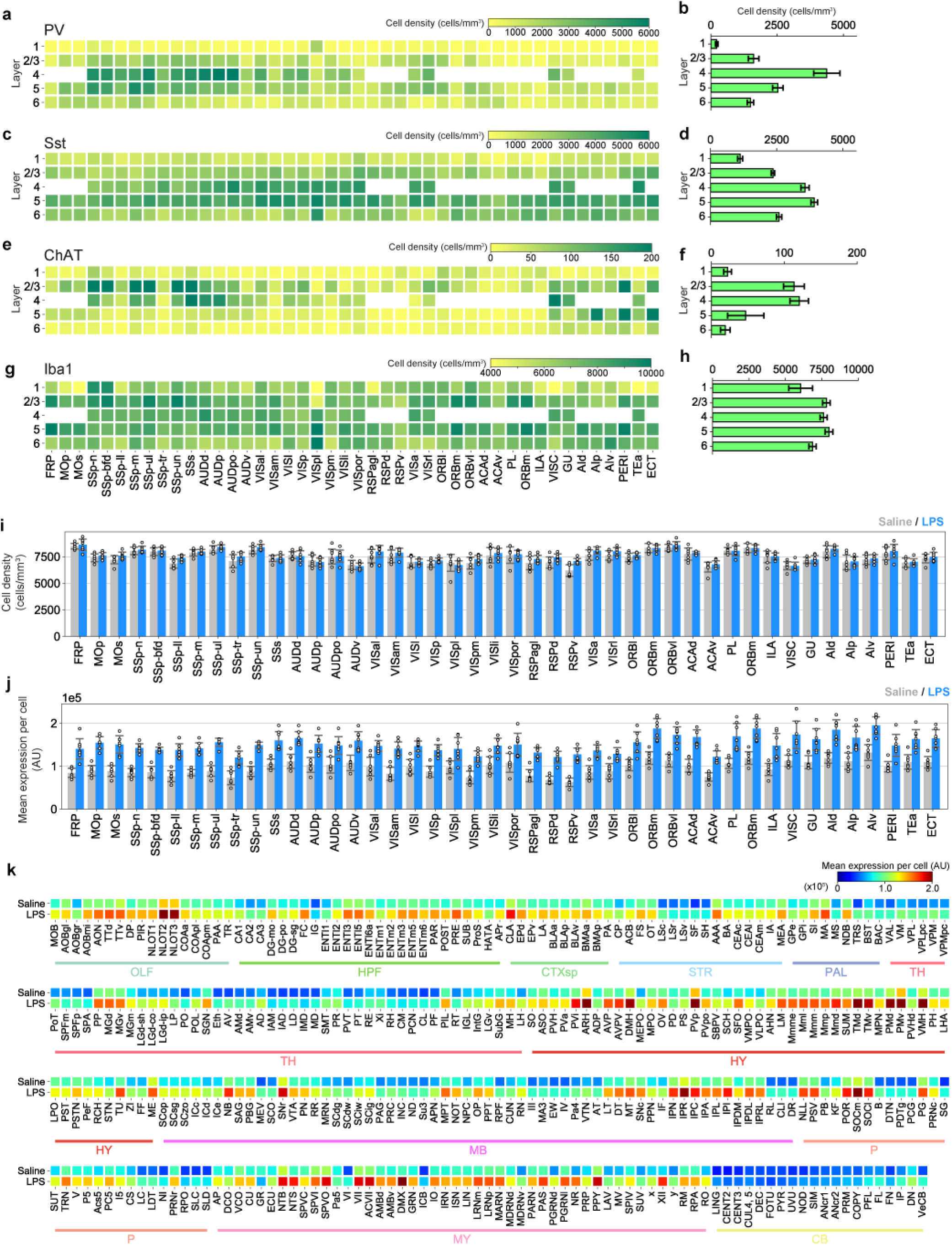
Whole-brain analysis of PV, SST, ChAT, TH and Iba1 Expressing Cells. **a**, Heatmap showing PV expressing cell density. **b**, Layer-wise average density of PV expressing cells. **c**, Heatmap showing SST expressing cell density. **d**, Layer-wise average density of SST expressing cells. **e**, Heatmap showing ChAT expressing cell density. **f**, Layer-wise average density of ChAT expressing cells. **g**, Heatmap showing Iba1 expressing cell density. **h**, Layer-wise average density of Iba1 expressing cells. **i**, Mean Iba1+ cell density in the isocortex. Saline (gray) and LPS (blue) groups are compared. Data are shown as mean ± STD (*n* = 7 for each group). **i**, Mean Iba1 expression level per cell in the isocortex. **j**, Heatmap showing mean Iba1 expression level per cell in all brain regions outside the isocortex, comparing saline and LPS groups. Brain region acronyms follow the ontology defined by the Allen Brain Atlas.

**Supplementary Figure 5.**
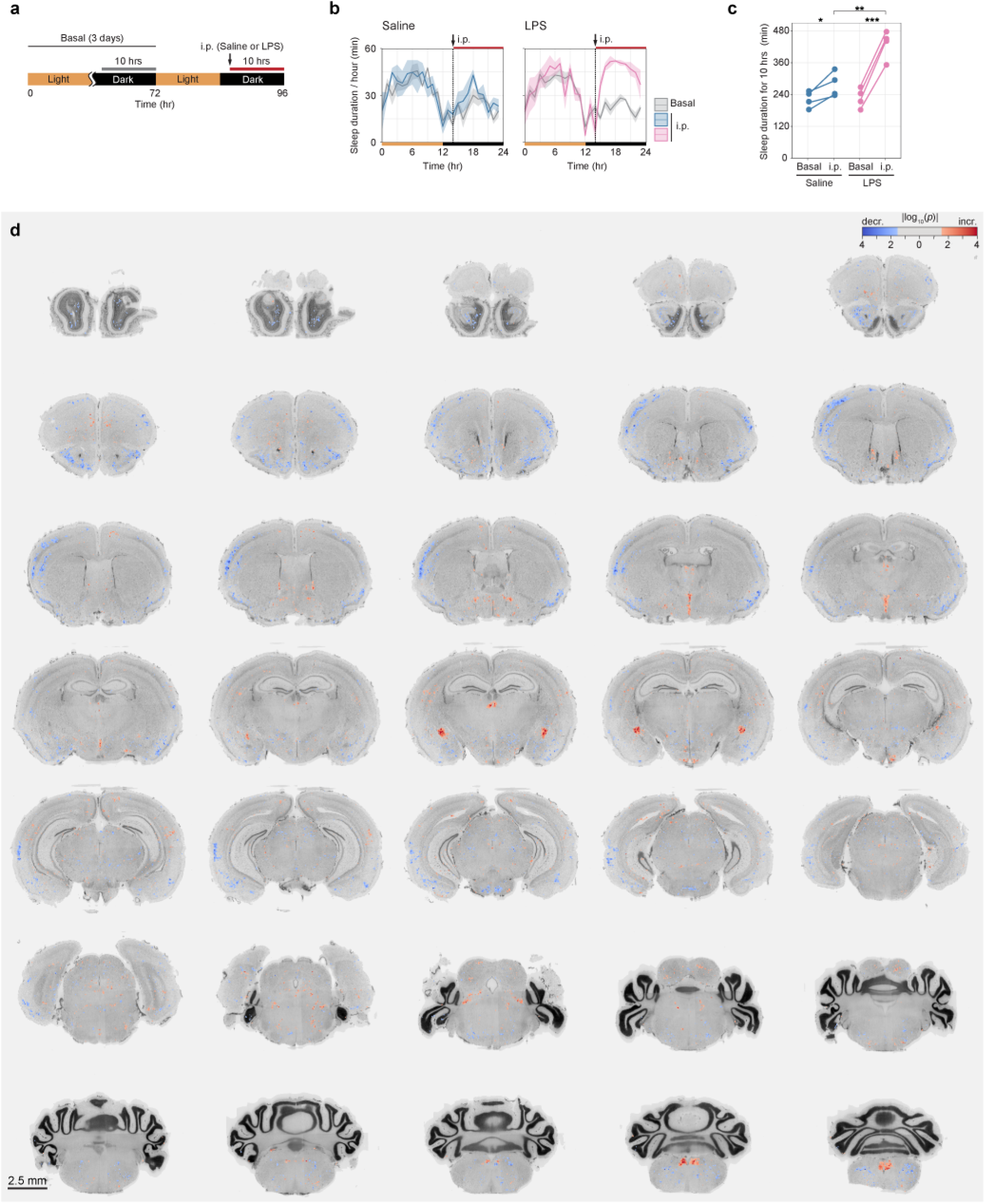
Whole-brain analysis of c-Fos expression under LPS administration. **d**, Voxel-wise p-value heatmap showing the affected regions by LPS. P-values of individual voxels were computed by c-Fos+ cell count between saline- and LPS-administered groups. The color lookup table is log scaled (base 10), where red color represents the regions that were activated (i.e. more c-Fos+ cells) by LPS, and blue represents the repressed regions. Voxels with no significance (*p* > 0.05) were uncolored. Background is nuclear staining image of CUBIC-Atlas for navigation. Step between consecutive slices is 0.34 mm.

**Supplementary Figure 6.**
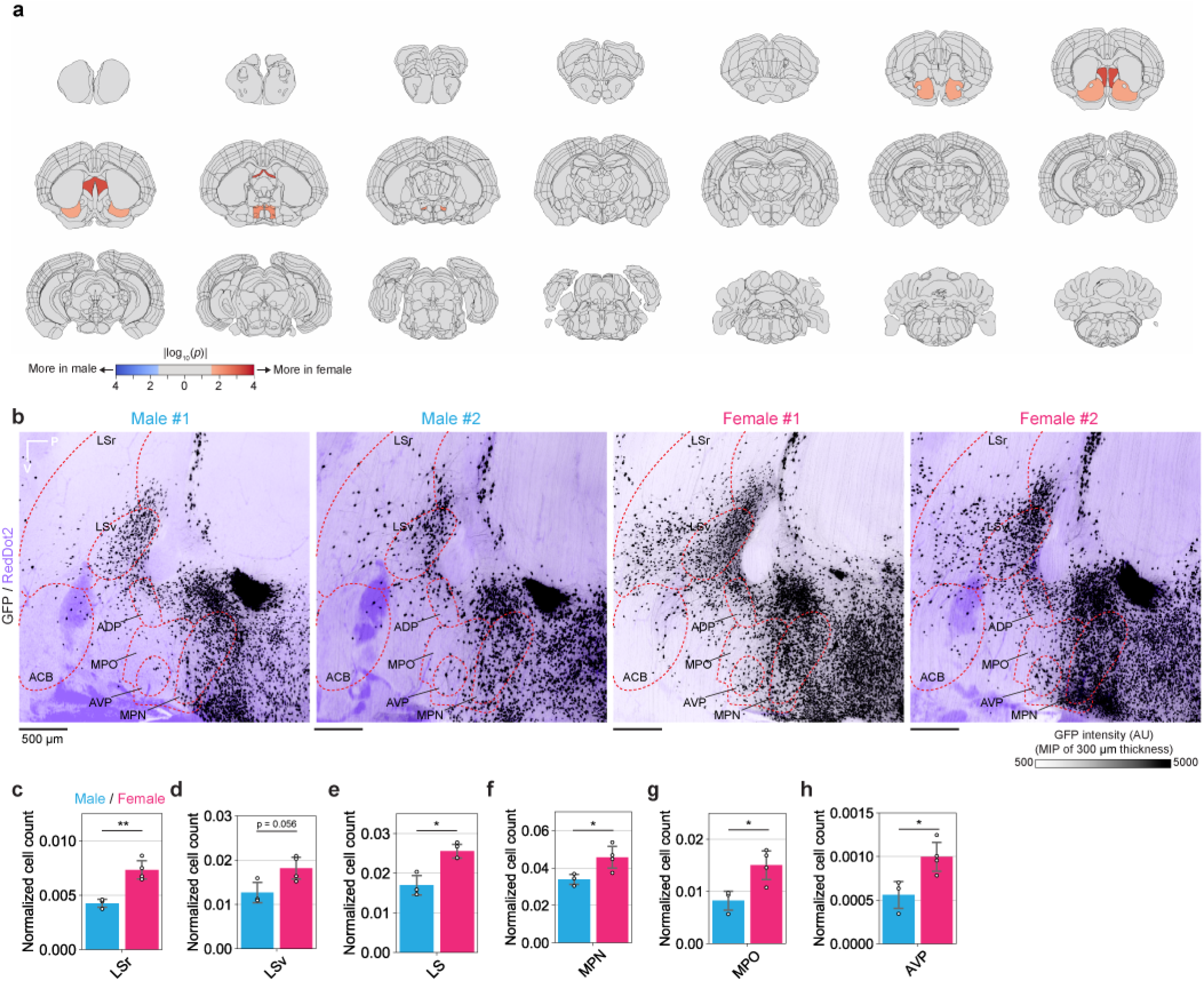
Additional Data of Rabies Virus Tracer Experiment. **a**, P-value heatmap where the number of input cells were compared between male and female brains. The color lookup table is log scaled (base 10), where red color represents the regions where more input cells were found in female brains, and blue represents the inverse. Regions with no statistical significance (*p* > 0.05) were assigned a gray color. **b**, Raw GFP (black) and nuclear staining (RedDot2, purple) images around lateral septal nucleus (LS) and medial prepoptic area (MPO). The images are digitally reconstructed sagittal sections. Maximum intensity project (MIP) spanning 300 μm thickness. **c-h**, The plot shows the normalized input cell count in regions where sexual dimorphisms were suggested. Of note, using RV injection and slice-based observation, [78] investigated the input cell population of pro-opiomelanocortin (POMC) neurons and agouti-related peptide (AgRP) neurons in the ARH, another dominant cell types in the ARH. The brain areas containing the input cells to ARH^*Kiss1+*^ neurons largely overlaped with those of POMC neurons and AgRP neurons. In some areas, however, interesting differences were observed. For example, in ventral tegmental nucleus (VTA) and nucleus incertus (NI), no input cells were detected for ARH^*Kiss1+*^, while some input cells were reported to exist for POMC and AgRP On the other hand, basomedial amygdalar nucleus (BMA), posterior amygdalar nucleus (PA), septofimbrial nucleus (SF), central amygdalar nucleus (CEA), substantia innominate (SI), triangular nucleus of septum (TRS), posterior intralaminar thalamic nucleus (PIL), midbrain reticular nucleus (MRN) and parabrachial nucleus (PB) contained small number of input cells to ARH^*Kiss1+*^ neurons (Fig. 5e,f,g), while they were not reported for POMC and AgRP neurons. This absence of input cells may reflect the actual biological differences, or it may reflect the superior sensitivity of our experimental methods to detect sparse populations. Further investigations are needed to draw conclusions. \**p* < 0.05, \*\**p* < 0.01; Welch’s t-test. Brain region acronyms follow the ontology defined by the Allen Brain Atlas.

Supplementary Table 2: Number of PV, SST, ChAT and Th expressing cells of adult mouse brains

The columns G to R list the number of PV+ cells (*n* = 4, data of each brain, the mean and standard deviations are shown) of all brain regions defined in Allen Brain Atlas (CCFv3, version 2017), along with the total fluorescence intensity. The columns S to AD list the results for SST+ cells (*n* = 4). The columns AE to AP list the results for ChAT+ cells (*n* = 4). The columns AQ to BB list the results for Th+ cells (*n* = 4).

Supplementary Table 3: Number and expression level of Iba1 expressing cells of LPS- or saline-administered brains

The columns G to X list the number of Iba1+ cells of saline-administered group (*n* = 7, data of each brain, the mean and standard deviations are shown) of all brain regions defined in Allen Brain Atlas (CCFv3, version 2017), along with the total fluorescence intensity. The columns Y to AP list the results for LPS-administered group (*n* = 7).

Supplementary Table 4: Number and volume of Aβ plaques of *App*^NL-G-F/NL-G-F^ mouse brain

The columns list the number of Aβ plaques (*n* = 4, data of each brain, the mean and standard deviations are shown) of all brain regions defined in Allen Brain Atlas (CCFv3, version 2017), along with the total plaque volume and fluorescence intensity.

Supplementary Table 5: Number and expression level of c-Fos expressing cells of LPS- or saline-administered brains

The columns G to R list the number of c-Fos+ cells of saline-administered group (*n* = 4, data of each brain, the mean and standard deviations are shown) of all brain regions defined in Allen Brain Atlas (CCFv3, version 2017), along with the total fluorescence intensity. The columns S to AD list the results for LPS-administered group (*n* = 4).

Supplementary Table 6: Number of input cells projecting to ARH^*Kiss1+*^ neurons

The columns list the number of input cells (i.e. GFP+ and mCherry-cells) (*n* = 3 for male, *n* = 4 for female, data of each brain, the mean and standard deviations are shown) of all brain regions defined in Allen Brain Atlas (CCFv3, version 2017), along with the total GFP intensity.

Supplementary Movie 1: Quick guide to CUBIC-Cloud

The movie shows the general usage of the CUBIC-Cloud.

